# The guanine nucleotide exchange factor Rin-like acts as a gatekeeper for T follicular helper cell differentiation via regulating CD28 signaling

**DOI:** 10.1101/2022.06.23.497284

**Authors:** Lisa Sandner, Marlis Alteneder, Ramona Rica, Barbara Woller, Eleonora Sala, Tobias Frey, Anela Tosevska, Ci Zhu, Moritz Madern, Pol Hoffmann, Alexandra Schebesta, Ichiro Taniuchi, Michael Bonelli, Klaus Schmetterer, Matteo Iannacone, Mirela Kuka, Wilfried Ellmeier, Shinya Sakaguchi, Ruth Herbst, Nicole Boucheron

## Abstract

T follicular helper (Tfh) cells are essential for the development of germinal center B cells and high-affinity antibody producing B-cells in human and mice. Here, we identify the guanine nucleotide exchange factor (GEF) Rin-like (Rinl) as a negative regulator of Tfh generation. Loss of Rinl leads to an increase of Tfh in aging, upon *in vivo* immunization and acute LCMV Armstrong infection in mice, and in human CD4^+^ T cell *in vitro* cultures. Further, adoptive transfer experiments using WT and Rinl-KO naïve CD4^+^ T cells unraveled T cell-intrinsic functions of Rinl. Mechanistically, Rinl regulates CD28 internalization and signaling, thereby shaping CD4^+^ T cell activation and differentiation. Thus, our results identify the GEF Rinl as a negative regulator of global Tfh differentiation in an immunological context and species-independent manner, and furthermore connect Rinl with CD28 internalization and signaling pathways in CD4^+^ T cells, demonstrating for the first-time the importance of endocytic processes for Tfh differentiation.

**Highlights:**

- Rinl-KO CD4^+^ T cells show increased Tfh differentiation in a context independent manner
- The regulation of Tfh differentiation is T cell-intrinsic
- Rinl controls CD28 endocytosis and shapes Tfh-specific CD28 signal transduction
- Human Tfh differentiation is regulated by Rinl

## Introduction

Small GTPases have been described as regulators of a variety of fundamental processes such as cell proliferation, differentiation or survival, with Rab GTPases primarily being involved in trafficking processes (Agola et al., 2011; Pfeffer, 2013). They act as so-called molecular switches binding to either GDP (inactive state) or GTP (active state). The change between the two conformational states is carried out by guanine nucleotide exchange factors (GEFs) and GTPase-activating proteins (GAPs) (Trabalzini and Walker, 2013). GEFs facilitate release of GDP, thereby enabling binding of GTP to the G protein (Bos et al., 2007). To this date, several families of GEFs for Rab GTPases have been described, with a large group comprising the Vps9-domain containing family where the Vps9-domain catalyzes the nucleotide exchange (Carney et al., 2006; Hama et al., 1999).

Ras-interaction/interference (Rin) proteins belong to this family and consist of 4 members called Rin1-3 and Rin-like (Rinl). Rin1-3 functions were elucidated in several *in vitro* and *in vivo* studies. They were described as GEFs for Rab5 sharing besides the Vps9 domain, also Src homology 2 (SH2), proline-rich (PR), RIN family homology (RH), and Ras association (RA) domains. The clinical outcome of deficiencies or overexpression in Rin1-3 range from neurological conditions like Alzheimer’s disease to macrocephaly, alopecia, cutis laxa, and scoliosis (MACS) syndrome (Kajiho et al., 2003; Saito et al., 2002; Tall et al., 2001). The latest member of Rin protein family, Rinl was originally identified as an interaction partner of the muscle specific receptor tyrosine kinase MuSK. The structure of Rinl highly resembles the structure of other Rin proteins, however, the Ras association domain is absent. Previous studies identified Rinl as GEF for Rab5 and Rab22 (Woller et al., 2011) and demonstrated its interaction with ankyrin-repeat and sterile-alpha motif (SAM) domain-containing (Anks) protein Odin, thereby facilitating EphA8 degradation (Kajiho et al., 2012). Interestingly, high Rinl expression was detected in lymphoid organs which indicates a role of Rinl in lymphoid cells. However, knowledge on the function of Rinl in primary cells was missing.

T follicular helper cells (Tfh) are a subset of CD4^+^ T helper cells and were first described to be present in human tonsils and blood (Kim et al., 2001; Schaerli et al., 2000). They are localized in Germinal Centers (GCs) and express high levels of CXC chemokine receptor 5 (CXCR5) while having low levels of CC chemokine receptor 7 (CCR7), thus being able to migrate to B-cell follicles to provide help for B-cells (Kim et al., 2001; Schaerli et al., 2000). Tfh are important players in the immune response against pathogens like bacteria or viruses and were also implicated in diseases including autoimmune diseases, immunodeficiencies, allergic asthma and lymphomas (Tangye et al., 2013). Moreover, they are important for the generation of high-affinity antibody responses after for example flu vaccines and SARS-CoV2 infections (Gong et al., 2014; Kaneko et al., 2020). Therefore, identifying factors that control Tfh differentiation and function is pivotal for the development of new treatments, diagnostic tools and vaccines.

Since the identification of B cell lymphoma (Bcl) 6 as major transcription factor (TF) of Tfh (Johnston et al., 2009; Nurieva et al., 2009; Yu et al., 2009), substantial progress was made in identifying factors involved in generation and function of these cells. The differentiation of Tfh was found to be a process in multiple stages facilitated by numerous factors. It was shown that CD28-deficient mice show reduced T helper (Th) 1 and Tfh cell expansion after infection or immunization (Shahinian et al., 1993; Walker et al., 1999). In addition, CD28 was reported to be important for the early stage of Tfh differentiation, while ICOS was identified as a key factor for sustaining Tfh differentiation (Weber et al., 2015).

Here, we use Rinl deficient mice to study the role of Rinl *in vivo*. Gene expression studies revealed high Rinl abundance in T cells. Further analysis of lymphoid organs and adoptive transfer experiments showed an increase of Tfh in absence of Rinl upon ageing, immunization and LCMV Armstrong infection. This universal regulation of Tfh differentiation under different immunological settings is due to T cell-intrinsic effects. Transcriptome analysis of naïve CD4^+^ T cells led to the identification of CD28 as upstream regulator being affected by loss of Rinl. We show that Rinl plays a role in the regulation of CD28 internalization which produced qualitative changes in CD28 downstream signaling *ex vivo* and shaped the differentiation into Tfh after CD28 induction *in vivo*.

Taken together, our findings suggest Rinl as a novel regulator of Tfh differentiation, highlighting the role of GEFs in T helper cell differentiation through involvement in trafficking of signaling receptors and subsequent changes in their signaling strength and/or quality.

## Results

### Rinl controls the homeostasis of CD4^+^ T helper cells in secondary lymphoid organs

The family of Rin proteins includes 4 members termed Rin1-3 and Rinl, all sharing the SH2, Vps9 and RH domain. The RA domain is present in Rin1-3 but absent in Rinl suggesting differences in function and/or localization of this GEF compared to the other family members **(Supplementary Fig. 1A)** (Woller et al., 2011). Previously performed tissue expression analyses reported highest Rinl expression in murine thymus, spleen and lymph node (Yue et al., 2014). Indeed, analysis of Rinl expression in WT mice showed the highest abundance of Rinl in lymphoid organs such as thymus, spleen, bone marrow (BM) and lymph nodes **(Supplementary Fig. 1B)** (Woller et al., 2011). To dissect the role of Rinl *in vivo*, Rinl-KO mice were generated by knock in (KI) of a STOP-IRES-LacZ::GFP cassette in the reading frame of exon 4 followed by floxed neomycin. Resulting *Rinl*^*+/KI-neo*^mice were crossed to CMV-Cre mice to delete the neomycin **(Supplementary Fig. 1C)**. The insertion led to an inactivation of the Rinl allele and absence of Rinl protein in Rinl^*KI/KI*^ mice as demonstrated by immunoblot assay but GFP or β-Gal were not detectable **(Supplementary Fig. 1D)**. The inactivated allele is henceforth referred to as a “knockout” allele (Rinl-KO). Rinl-KO mice were developmentally normal and fertile being born at normal Mendelian frequencies. Due to high expression of Rinl in lymphoid organs, we first aimed to study T cell development in the thymus of WT and Rinl-KO mice. Frequencies of DN, DP and CD4^+^/CD8^+^ SP subsets were comparable between Rinl deficient and control mice indicating Rinl to be dispensable for the development of T cells **(Supplementary Fig. 1E,F)**. As Rinl showed high expression in splenic CD4^+^ and CD8^+^ T cells **(Supplementary Fig. 1G)** and also an increase in gene expression in CD4^+^ effector compared to naïve CD4^+^ T cells **(Supplementary Fig. 1H)**, we analyzed lymphocyte homeostasis in the spleen **(Supplementary Fig. 1I)**. We detected a significant relative increase in frequencies of B cells and effector CD4^+^ (CD4^+^CD44^+^) T cells in mice aged between 8-12 weeks. Furthermore, Rinl-KO mice showed a strong increase in total splenocytes with consequent significantly increased absolute numbers of B and T lymphocytes, in particular effector CD4^+^ (CD4^+^CD44^+^) T cells, whereas CD8^+^ T lymphocytes were unaltered **(Supplementary Fig. 1I**,**J)**.

Strikingly, aged Rinl-KO mice (36-40weeks) showed splenomegaly, increased spleen weight and increased splenocyte numbers **(Fig. 1A,B)**. They accumulated more T cells, CD4^+^ and effector CD4^+^ T cells (CD62L^−^CD44^+^) compared to WT mice. Additionally, an increase of total Tfh and GC B cells was detected **(Fig. 1C, For gating strategy please refer to Supp Fig. 2A)**, while other T helper subsets were not altered **(Supplementary Fig. 2B**,**C)**. In lymph nodes of aged mice, an increase in CD4^+^ and Tfh cell frequencies was also observed in absence of Rinl **(Supplementary Fig. 2D)** indicating organ-dependent alterations in lymphocyte subsets. Altogether, our immunophenotyping of Rinl mice suggest a regulatory role of Rinl in CD4^+^ T helper cell activation/differentiation, in particular towards Tfh.

**Figure 1:**
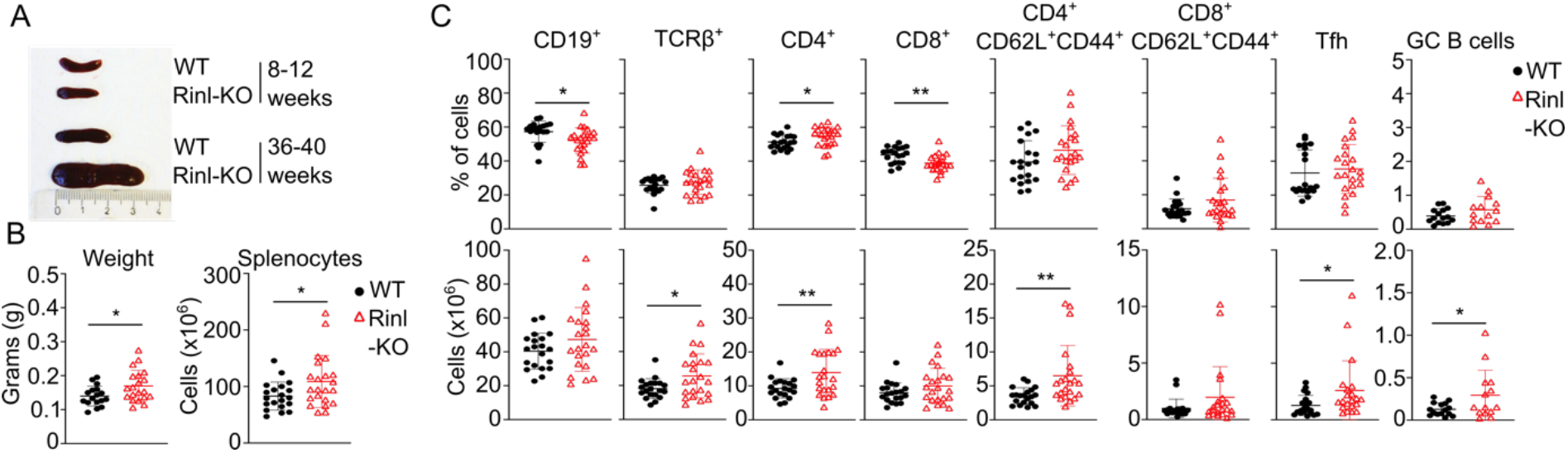
Absence of Rinl leads to alterations in immune homeostasis in older mice. **(A)** Representative picture of spleens of WT and Rinl-KO mice of 8-12 and 36-40 weeks of age. **(B)** Diagrams show weight and total leukocyte count of splenocytes of 36-40 week old WT and Rinl-KO mice. **(C)** Summary graphs show the percentages (upper panel) and cell counts (lower panel) of major lymphocyte subsets analyzed. The summary of 17-20 mice per genotype analyzed in 4 independent experiments is shown. Data were statistically analyzed using unpaired two-tailed t-tests **(B**,**C)**. *p < 0.05, **p<0.01.

### Rinl regulates Tfh differentiation and GC B cell generation in homeostasis and upon immunization

To define the role of Rinl in CD4^+^ T helper (Th) cells in more detail, we first investigated the differentiation potential of WT and Rinl-KO naïve CD4^+^ T cells into Th1, Th2, Th17 and Treg cells *in vitro*. Frequencies of key cytokine-producing or lineage-specific transcription factor expressing cells of all major Th subsets were comparable suggesting Rinl to be dispensable for regulating differentiation of those T helper subsets *in vitro* (**Supplementary Fig 3A**,**B**). First analyses of Tfh (defined by CXCR5 and PD-1) and GC B cells (defined by GL7 and CD95) were performed in Peyer’s Patches (PP), a lymphoid structure favoring the generation of those cells due to constant stimulation by external factors like the microbiome (Jones et al., 2016). In these specimen, Rinl-KO mice showed a significant increase of both cell subsets under homeostatic conditions **(Fig. 2A,B)**. We furthermore demonstrated a high expression of Rinl in Tfh while expression in GC B cells was low **(Fig. 2C**) suggesting a specific role of Rinl in the generation of Tfh.

**Figure 2:**
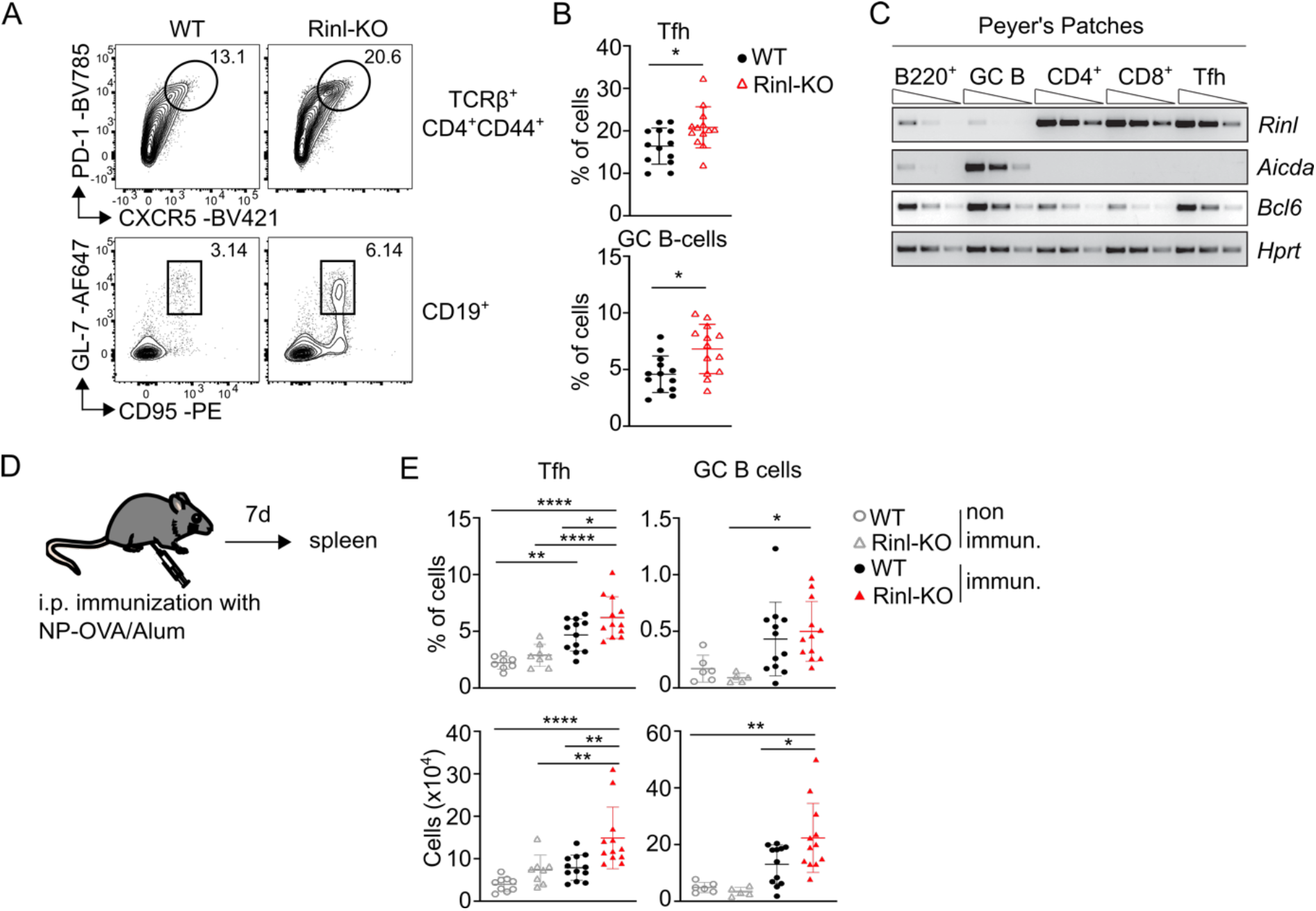
Rinl controls Tfh and GC B cell generation in homeostasis and upon immunization. **(A)** Representative contour plots show Tfh and GC B cells in Peyer’s patches of WT or Rinl-KO mice. Numbers in plots indicate percentages of cells. **(B)** Quantification of **(A). (C)** Semi-quantitative PCR of *Rinl, Aid and Bcl6* expression in different cell subsets in Peyer’s Patches of WT mice. *Hprt* expression was used as control. **(D)** Experimental scheme of immunization experiment. **(E)** Summary graphs depict percentages (upper panel) and cell numbers (lower panel) of Tfh and GC B cells recovered from spleens from non-immunized and immunized WT and Rinl-KO mice. Data show a summary of 11-13 **(B)** and 5-8 **(E)** mice per group analyzed in 3-4 independent experiments **(B**,**E)**. Data were statistically analyzed using unpaired two-tailed t-tests **(B)** or 1-way ANOVA analysis followed by Tukey’s multiple-comparisons test **(E)**. *p < 0.05, **p< 0.01, and ***p < 0.001, ****P<0.0001.

Next, we sought to determine the importance of Rinl during Tfh differentiation following immunization. We immunized WT and Rinl-KO mice intraperitoneally (i.p) with 100 μg/mL 4-Hydroxy-3-nitrophenylacetyl hapten conjugated to Ovalbumin (NP-OVA) in the Th2-driving Imject™ Alum Adjuvant (Alum) **(Fig. 2D)**. As expected, immunization with NP-OVA/Alum led to an increase in Tfh frequencies and cell numbers compared to non-immunized control mice 7 days post immunization. Strikingly, we detected an increase in Tfh in relative and absolute numbers in Rinl-KO mice compared to WT mice **(Fig. 2E)** along with an increase in GC B cells. Immunization was also performed using Complete Freund’s Adjuvant (CFA)**(Supplementary Fig.3C)**, which augments mostly Th1- and Th17-mediated responses (Ciraci et al., 2016; Yang and Hayglass, 1993). Similar to results gained with Alum, Tfh were significantly increased in Rinl-KO mice in comparison to WT counterparts **(Supplementary Fig.3D)**, whereas other T helper subsets were unaltered **(Supplementary Fig.3E)**. To conclude, our data show evidence that Rinl specifically controls the differentiation of Tfh *in vivo*.

### Rinl is a negative regulator of Tfh differentiation upon LCMV infection

We next aimed to study the effect of Rinl on Tfh differentiation in the context of an infection setting. We used an acute infection model with Lymphocytic choriomeningitis virus (LCMV) which induces the generation of virus specific Th1 and Tfh cells, in addition to a CD8^+^ T cell response (Hale et al., 2013; Marshall et al., 2011). At day 8 after infection with LCMV, Rinl-KO mice showed no change in frequencies of lymphocyte subsets (**Fig. 3A,** for gating strategy please refer to **Supplementary Figure 4A, Supplementary Fig. 4B,)**. However, a significant increase in splenocyte numbers was observed, along with a prominent expansion of total CD4^+^ T helper cells in Rinl-KO compared to WT mice **(Fig. 3B)**. Among CD4^+^ T cells a significant increase of effector CD4^+^ (CD4^+^CD44^+^) and Tfh (PD-1^+^CXCR5^+^) was observed in absence of Rinl. Moreover, while a similar amount of virus-specific gp66 tetramer-positive CD4^+^ T cells in Rinl-KO and control mice was detected, a higher number of gp66 tetramer-positive PD-1^+^CXCR5^+^ Tfh were present in Rinl-KO mice **(Fig. 3B)**. The number of gp66 tetramer-positive PSGL1^+^Ly6C^+^ Th1 was unaltered, indicating a suppressive role of Rinl selectively on Tfh differentiation during viral infection **(Fig. 3B)**(Xu et al., 2015). Similar to immunization experiments, LCMV infection led to an increase in B cells 8 days post infection (dpi) and a significant enhancement of GC B cells and IgG1^+^ B cells in absence of Rinl at day 21 after infection **(Fig. 3C,D** and **Supplementary Fig. 4C)**. Additionally, IgG1 antibodies in serum of Rinl-KO mice 21dpi showed a tendency of increasement **(Fig. 3C)**. Overall, infection experiments disclosed a role of Rinl in virus-specific Tfh differentiation and generation of virus-specific antibodies.

**Figure 3:**
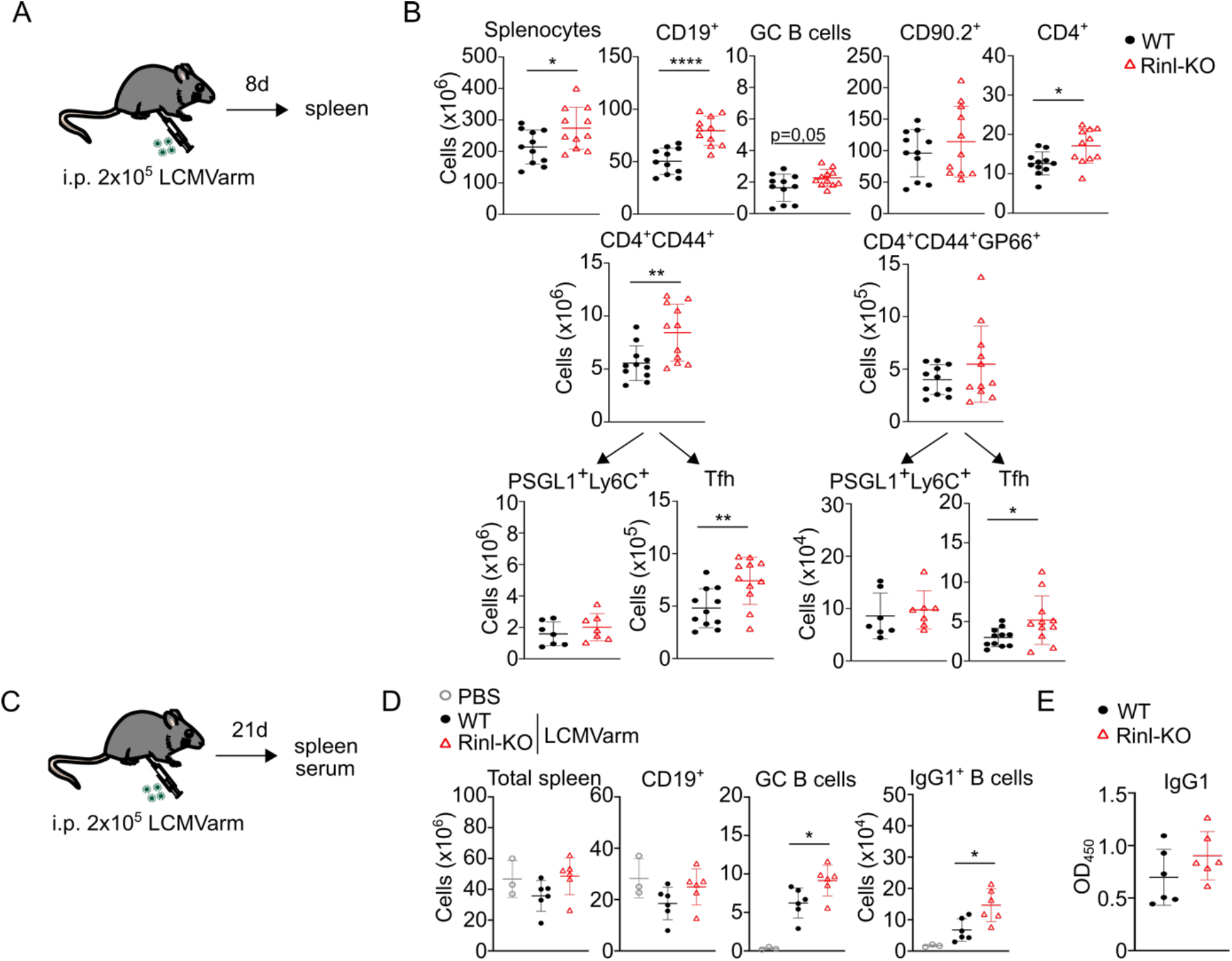
Rinl regulates Tfh and B cell response upon LCMV Armstrong infection. **(A)** Experimental scheme. **(B)** Summary diagrams depict cell numbers of a variety of lymphocyte subsets from spleens of WT and Rinl-KO mice 8 days post infection. **(C)** Experimental scheme of LCMV Armstrong infection to study B cell responses. **(D)** Summary diagrams depict cell numbers of B cell subsets from spleens of WT and Rinl-KO mice 21 days post infection **(E)** Virus-specific IgG1 levels in serum 21 days post infection. Data show a summary of 11 **(B)** or 6 **(D**,**E)** mice per group analyzed in 3 **(B)** or 2 **(D**,**E)** independent experiments. Data were statistically analyzed using unpaired two-tailed t-tests. *p < 0.05, **p < 0.01, ****p<0.0001

### Rinl has a T cell-intrinsic effect on T cell activation on early Tfh commitment without affecting overall CD4^+^ T cell activation, proliferation and migration/localization

To examine the intrinsic role of Rinl in early T cell activation and proliferation *in vivo*, we crossed WT and Rinl-KO mice to Ovalbumin (OVA) TCR transgenic mice further named as either WT OT-II^+^ or Rinl-KO OT-II^+^ mice. We subsequently performed adoptive transfer experiments in which we isolated WT or Rinl-KO OT-II^+^ naïve CD4^+^ CD45.2^+^ T cells, labelled them with CFSE and injected them intravenously (i.v) into CD45.1^+^CD45.2^+^ recipient mice. Two days after immunization, transferred (CD45.2^+^CD45.1^−^) CD4^+^ T cells in draining (popliteal) lymph nodes (dLN) and non-draining (axillary) lymph nodes (ndLN) were analyzed for proliferation and early activation marker expression **(Fig. 4A and Supplementary Fig. 5A)**. We did not observe differences in frequencies of transferred cells, proliferation (defined as CFSE^low^), and activation measured via CD25 and CD69 expression **(Supplementary Fig. 5B)**. However, Rinl-KO OT-II^+^ cells expressed significantly higher amounts of CD44^+^ fraction and the important Tfh markers CXCR5 and ICOS **(Fig. 4B,C)** (Choi et al., 2011). We furthermore performed immunohistochemistry of dLNs at day 3 and day 5 after immunization to define localization of transferred WT or Rinl-KO CD4^+^CD45.2^+^ T cells. At day 3 after immunization, we detected antigen specific CD45.2^+^ T cells of both genotypes in the T cell area close to the T-B cell border. Some of the T cells then migrate further into the B cell follicles at day 5 suggesting proper migration and localization of T cells also in absence of Rinl during early activation **(Supplementary Fig. 5C)**. To conclude, our data demonstrate a role of Rinl in early commitment towards Tfh but not in overall T cell activation, proliferation and migration.

**Figure 4:**
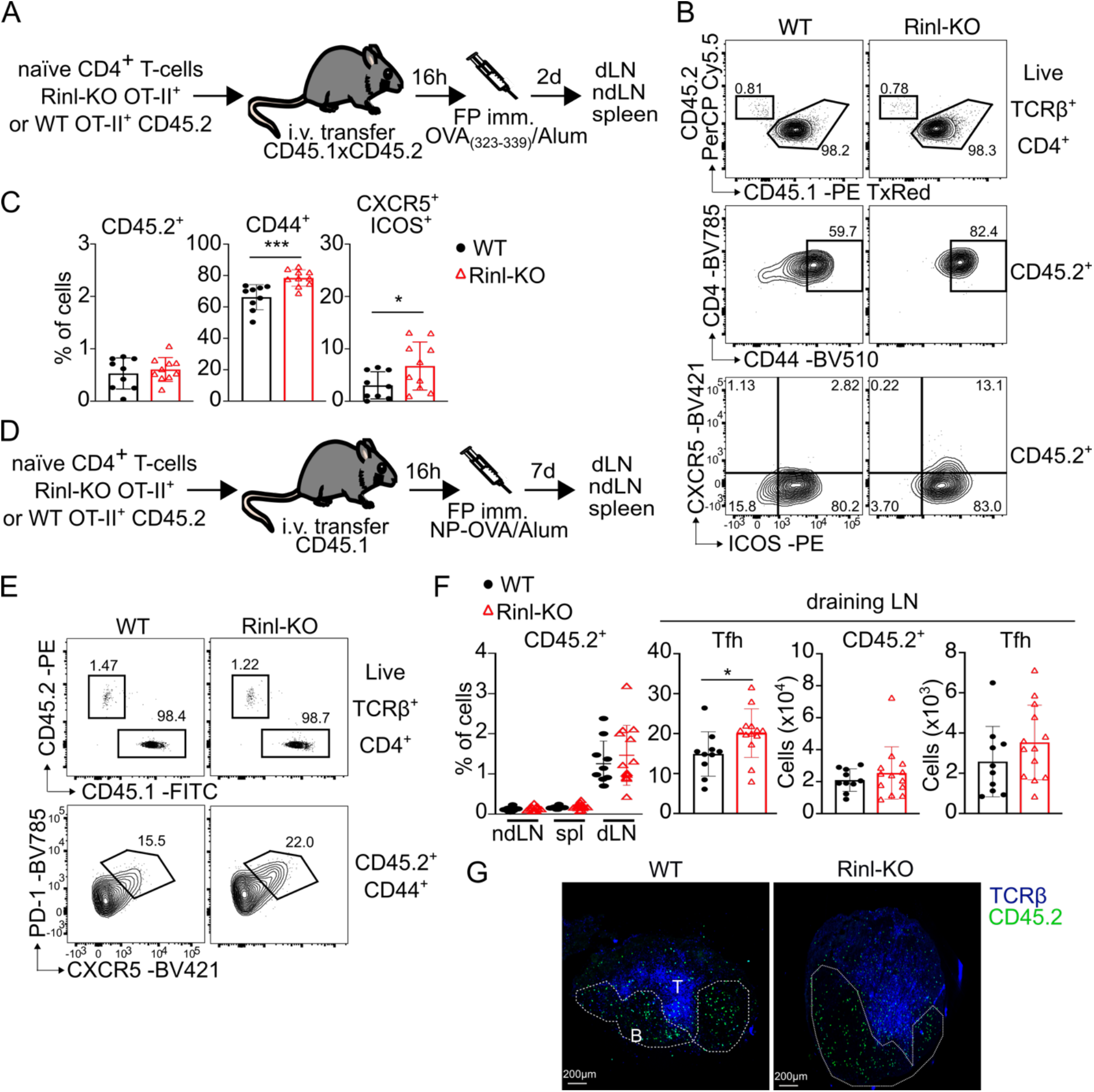
Increased Tfh differentiation in Rinl-KO mice is T cell intrinsic. **(A)** Experimental scheme. **(B)** Representative contour plots of dLN samples taken from WT and Rinl-KO mice showing transferred cells (CD45.2^+^), effector cells (CD44^+^) and pre-Tfh (ICOS^+^CXCR5^+^). **(C)** Summary of **(B). (D)** Experimental scheme. **(E)** Representative contour plots of dLN samples taken from WT and Rinl-KO mice showing transferred cells (CD45.2^+^) and Tfh (PD-1^+^CXCR5^+^). **(F)** Summary of **(E)**. Representative confocal images of dLN collected 7 days after immunization. Dashed lines represent edges of B cell follicles and were set based on CD21/35 staining. TCRß^+^ cells are shown in blue and Ag-specific CD45.2^+^ cells in green. Images are representative of at least 3-5 LNs from 2 independent experiments **(E)**. Data show a summary of 9-10 **(C)** and 10-13 **(F)** mice per group analyzed in 2 **(C)** or 3 **(F)** independent experiments. Data were statistically analyzed using unpaired two-tailed t-tests. *p < 0.05, ***p < 0.001.

### Rinl has a T cell-intrinsic effect on Tfh differentiation without affecting T cell homing

To study the differentiation potential of WT or Rinl-KO OT-II^+^ CD4^+^ T cells into full GC Tfh, we injected WT or Rinl-KO OT-II^+^ naïve CD4^+^CD45.2^+^ T cells into CD45.1^+^ recipient mice and performed FP immunization with NP-OVA/Alum the next day **(Fig. 4D)**. On day 7 after immunization, we first assessed migration of T cells in dLNs, spleens and ndLNs. Transferred (CD45.2^+^CD45.1^−^) CD4^+^ T cells were almost exclusively found in dLNs. Furthermore, frequencies in dLNs were unaltered between WT and Rinl-KO OT-II^+^ T cells indicating no impact of Rinl on the homing capacity of cells to secondary lymphoid organs **(Fig. 4E, F)**. Strikingly, we detected significant enhancement of PD-1^+^CXCR5^+^ Tfh among Rinl-KO OT-II^+^ T cells compared to WT OT-II^+^ T cells indicating a T cell-intrinsic expansion of Tfh upon loss of Rinl **(Fig. 4E, F)**. By performing immunohistochemistry of dLNs we further evaluated that both WT and Rinl-KO T cells are capable to reside within the B cell follicles and started to form clusters **(Fig. 4G)**.

Next, we investigated possible trans-effects due to soluble mediators released by CD4^+^ T cells by co-transferring WT OT-II^+^ naïve CD4^+^CD45.1^+^ T cells and either WT or Rinl-KO OT-II^+^ naïve CD4^+^CD45.2^+^ T cells i.v into TCRα^−/-^ mice **(Fig. 5A)**. Same frequencies of TCRβ^+^ cells and Tfh were detected in mice co-transferred with WT OT-II^+^CD45.1^+^/WT OT-II^+^CD45.2^+^ and WT CD45.1^+^/Rinl-KO CD45.2^+^ T cells 7 days post immunization **(Fig. 5B, C)**. Among Tfh cells, Rinl-KO CD45.2^+^ cells were significantly increased compared to WT CD45.2^+^cells co-transferred with WT CD45.1^+^ cells, showing an enhanced T cell-intrinsic Tfh differentiation potential of Rinl deficient CD45.2^+^ cells. In contrast, no difference in CD45.1 and CD45.2 proportion was detected in the non Tfh (PD-1^−^CXCR5^−^) population suggesting no influence of soluble factors of Rinl-KO T cells on WT T-cells favoring Tfh differentiation **(Fig. 5B, C)**.

**Figure 5:**
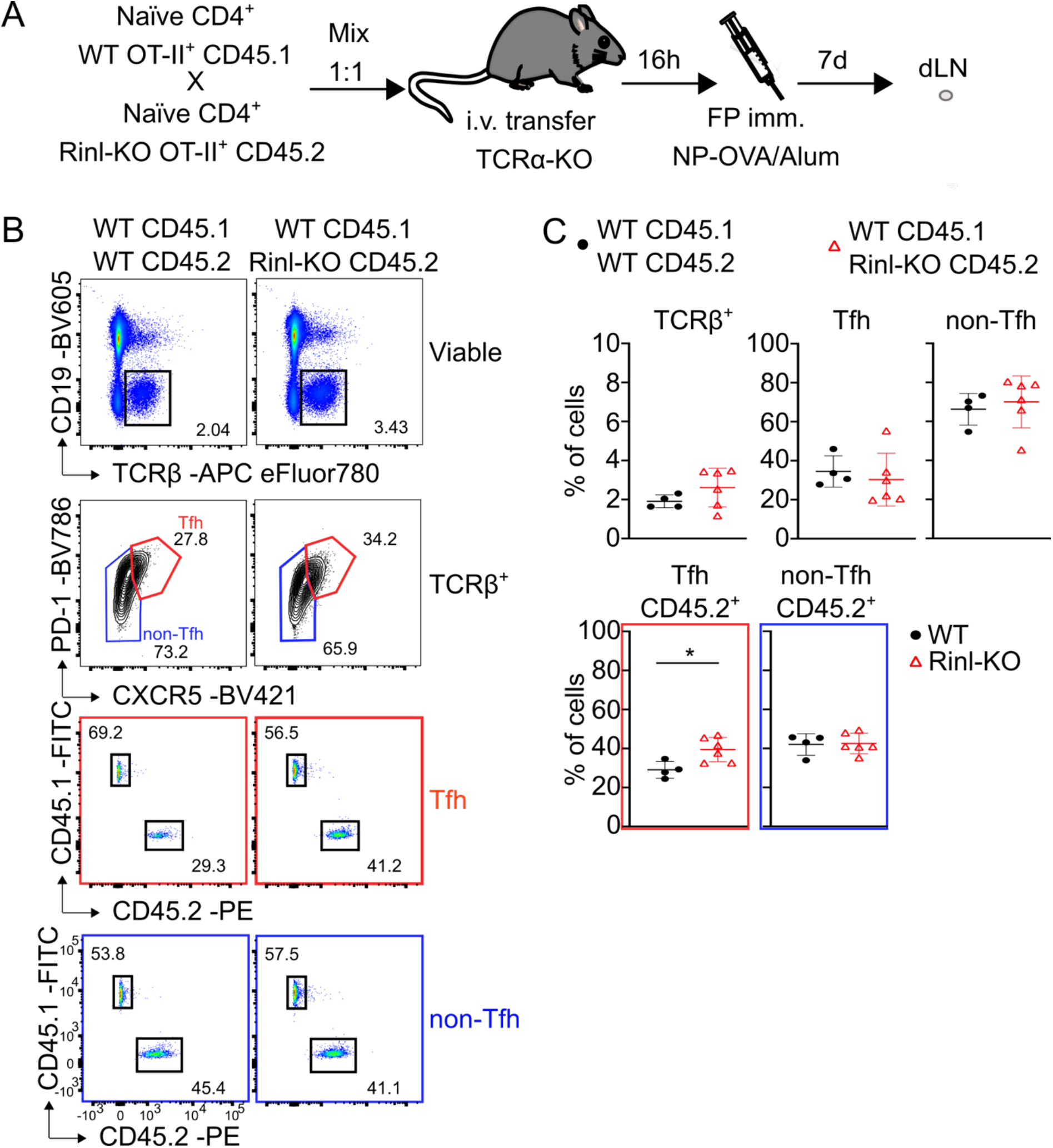
T cell trans effects do not affect Tfh differentiation in absence of Rinl. **(A)** Experimental scheme. **(B)** Representative flow cytometry plots of TCRβ^+^ cells, Tfh/non-Tfh and CD45.1/CD45.2 frequencies among Tfh and non-Tfh. Numbers indicate percentage of cells in the respective gate **(C)** Summary of **(B)**. Data show a summary 4-6 mice per group analyzed in 2 independent experiments. Data were statistically analyzed using unpaired two-tailed t-tests. *p < 0.05.

To exclude the involvement of non-T cells like DCs or B cells in Tfh differentiation in absence of Rinl, we crossed Rinl-KO mice to TCRα KO mice which lack all αβT cells further named Rinl-KO TCRα-KO mice. We then performed adoptive transfer experiments of WT OT-II^+^ naïve CD4^+^ T cells into either TCRα-KO or Rinl-KO TCRα-KO mice **(Supplementary Fig. 5D)**. 7 days post immunization comparable frequencies and cell counts of T cells and Tfh were detected in both recipient mice indicating no significant effect of Rinl-KO non-T cells on Tfh differentiation **(Supplementary Fig. 5E)**.

Collectively, adoptive transfer experiments indicate a T cell-intrinsic regulation of Tfh differentiation by Rinl which is not the consequence of an increase in secreted factors. This is also not due to changes in T cell homing or follicle formation but is the result of enhanced commitment towards this T helper subset in absence of Rinl. In addition, our data strongly suggest that other immune cells are not contributing to the enhanced Tfh differentiation when Rinl is not present.

### Rinl is a regulator of CD28 signaling pathway

In order to elucidate the molecular mechanisms by which Rinl regulates Tfh differentiation, we performed a transcriptome analysis of *ex vivo* sorted naïve CD4^+^ T cells (CD4^+^CD62L^+^CD44^−^) from spleens of WT and Rinl-KO mice using RNA sequencing (RNA-seq) approaches. Globally, 123 genes were down- and 217 genes upregulated in Rinl-KO naïve CD4^+^ T cells compared to WT cells **(Fig. 6A)**. Importantly, Ingenuity Pathway Analysis identified CD28 as a dysregulated upstream regulator **(Fig. 6B)**. CD28 was previously demonstrated to be indispensable for Tfh and GC formation (Ferguson et al 1996, Weber et al 2015) and fine-tuning of CD28 signaling was recently described as crucial factor in Tfh differentiation (Wan et al., 2021). To study this in more detail, we first analyzed CD28 expression on naïve and effector CD4^+^ T cells which was comparable between WT and Rinl-KO **(Supplementary Fig 6A)**.

**Figure 6:**
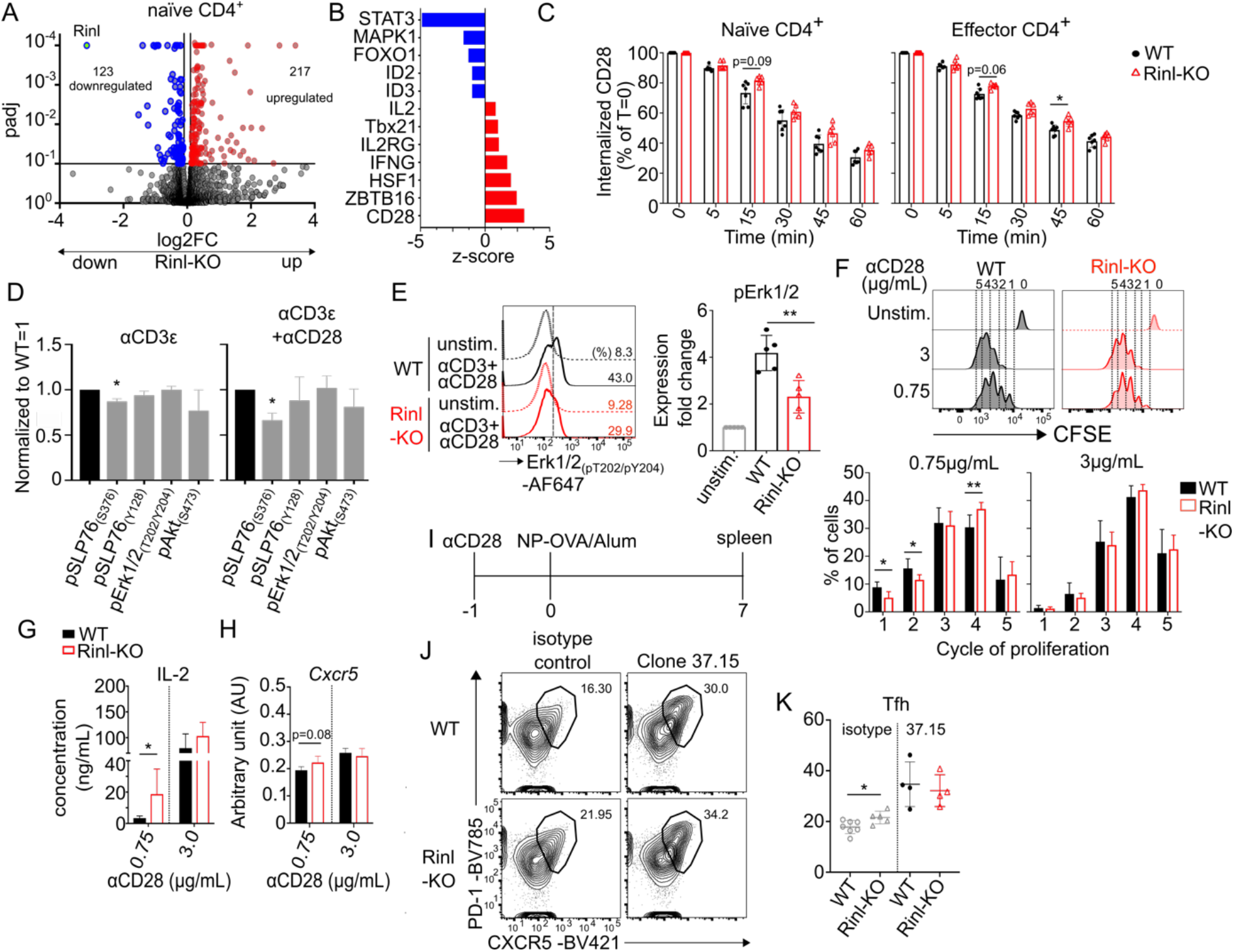
CD28 trafficking and signaling pathway is regulated by Rinl. RNA was isolated from sorted WT and Rinl-KO naïve CD4^+^ T cells (CD62L^+^CD44^−^) and subjected to RNA sequencing. **(A)** Volcano plot depicts the comparison of global gene expression profiles between WT and Rinl-KO naïve CD4^+^ T cells. The y-axis indicates adjusted p-values (-log10), the x-axis shows the log2 fold change. **(B)** Diagram shows top dysregulated upstream regulators identified by Ingenuity Pathway Analysis (Qiagen Inc). Regulators are plotted on y-axis and are predicted to be up-(positive z-score, red) or downregulated (negative z-score, blue) in Rinl-KO naïve CD4^+^ T cells. **(C)** Summary of CD28 internalization kinetic in naïve and effector CD4^+^ T cells. Internalization was calculated as percentage (%) of CD28 expression from time point 0 (T=0). **(D)** Summary of phosphorylation of signaling molecules 2 minutes after αCD3ε or αCD3ε+αCD28 stimulation of CD4^+^ T cells. **(E)** Representative histograms of Erk1/2_(pT202/pY204)_ of WT and Rinl-KO CD4^+^ T cells activated with immobilized αCD3+αCD28 for 48 hours are shown. Summary is depicted below. **(F)** Representative histograms of proliferation after activation of naïve CD4^+^ T cells with low (0,75 μg/mL) and high (3 μg/mL) αCD28 for 3 days. Summary of data is shown alongside. **(G)** Summary of IL-2 measured from supernatants of cells cultured as in **(F). (H)** Expression of *Cxcr5* was determined by RT-qPCR in naïve CD4^+^ T cells stimulated as described in **(F). (I)** Experimental setup. **(J)** Representative contour plots of Tfh of WT and Rinl-KO mice treated with isotype control or αCD28 (Clone 37.15) one day before immunization. **(K)** Summary of **(J)**. Data show a summary of 6-7 **(C)**, 3-4 **(D)**, 5 **(E)** 6 **(F**,**G)**, 4 **(H)** and 4-7**(J**,**K)** mice per group analyzed in 3-4 **(C**,**D, F, G, H)**, 5 **(E)** or 2 **(J**,**K)** independent experiments. Data were statistically analyzed using two-way ANOVA using Tukey multiple comparison’s test **(C)**, paired two-tailed **(E)** or unpaired two-tailed t-tests **(F**,**K)**. For the comparison of the relative geoMFI values between WT (set as 1 for each phospho protein) and Rinl-KO, a one-sample t-test was performed **(D)**. *p < 0.05, **p < 0.01.

CD28 expression is regulated by constant turnover of the receptor. As Rinl is a GEF for Rab5 GTPases involved in receptor trafficking, we examined its role in CD28 endocytosis and downstream signaling such as phosphorylation of SLP76, Akt or Erk1/2. Interestingly, endocytosis assay showed that the internalization of CD28 was decreased in Rinl-KO effector CD4^+^ T cells (CD4^+^CD44^+^) **(Fig. 6C)**. Moreover, combined αCD3ε and αCD28 stimulation of CD4^+^ T cells led to strongly reduced pSLP76 (Ser376) which was described to negatively regulate T cell activation (Di Bartolo et al., 2007) with unchanged Akt and Erk1/2 phosphorylation **(Fig. 6D)**. Intriguingly, after stimulation of cells with TCR and CD28 in culture for 48 hours, we detected reduced levels of Erk1/2 phosphorylation in Rinl-KO cells **(Fig 6E)**. The Erk pathway was recently described as repressive factor of Tfh differentiation and its sequential requirement during this process has been highlighted (Wan et al 2021). These data combined suggest in combination an effect of Rinl in shaping CD28 signaling quality.

To test whether CD28 signaling strength has an effect in absence of Rinl, we cultured naïve CD4^+^ T cells with constant αCD3ε and low and high αCD28 concentrations for 3 days and checked for proliferation, IL-2 production and *Cxcr5* gene expression **(Supplementary Fig 6B)**. Interestingly, we detected a significant increase in proliferation and IL-2 production in Rinl-KO cells stimulated with lower αCD28 concentration compared to WT cells **(Fig. 6F, G)**.

Proliferation in Rinl-KO T cells was also enhanced in a more physiological model when activation was performed with bone-marrow derived dendritic cells (BMDCs) in presence of CTL4-Ig **(Supplementary Fig 6C)**. Additionally, *Cxcr5* gene expression showed a tendency of increase in Rinl-KO cells activated with lower αCD28 concentration as detected by RT-PCR, whereas no difference between genotypes was detected with higher αCD28 concentrations **(Fig. 6H)**. Collectively, these data suggest a role of Rinl in fine-tuning CD28 downstream events, which indicate early Tfh commitment.

We next aimed to study the effect of Rinl on CD28 signaling and subsequent Tfh differentiation *in vivo*. Therefore, we treated WT or Rinl-KO mice with αCD28 or isotype control one day before i.p. immunization with NP-OVA Alum. Tfh cell frequency was analyzed on day 7 after immunization **(Fig. 6I)**. In agreement with a published study (Vogelzang et al., 2008), CD28 induction led to an increase of Tfh differentiation in both genotypes. Importantly, αCD28 treatment leads to similar frequencies of Tfh between the genotypes in contrast to isotype treatment where we could detect the increased differentiation into Tfh in spleens of Rinl-KO mice **(Fig. 6J)**. Of note, Tfr (FoxP3^+^ Tfh) frequencies are not affected by αCD28 treatment **(Supplementary Fig 6D)**. Altogether this suggest that Rinl is a negative regulator of Tfh by dampening the effect of CD28 stimulation on CD4^+^ T cells as strong CD28 signaling results in comparable Tfh in WT and Rinl-KO mice.

### Human Tfh differentiation is affected by Rinl

As high Rinl expression was also detected in human lymphoid tissues, we next aimed to study its role in human Tfh (Fagerberg et al., 2014). In contrast to murine cells, an *in vitro* differentiation protocol for human Tfh has been established (Locci et al., 2016). We isolated CD4^+^ T cells from human peripheral blood of healthy individuals and performed Crispr-Cas9 knockout of Rinl or control gRNA. After successful knockout we cultured cells under Tfh-skewing conditions. In accordance with the data from the murine experiments, Rinl-KO cells showed a higher differentiation potential into Tfh *in vitro* suggesting Rinl as negative regulator of Tfh differentiation also in humans **(Fig. 7A, B)**.

**Figure 7:**
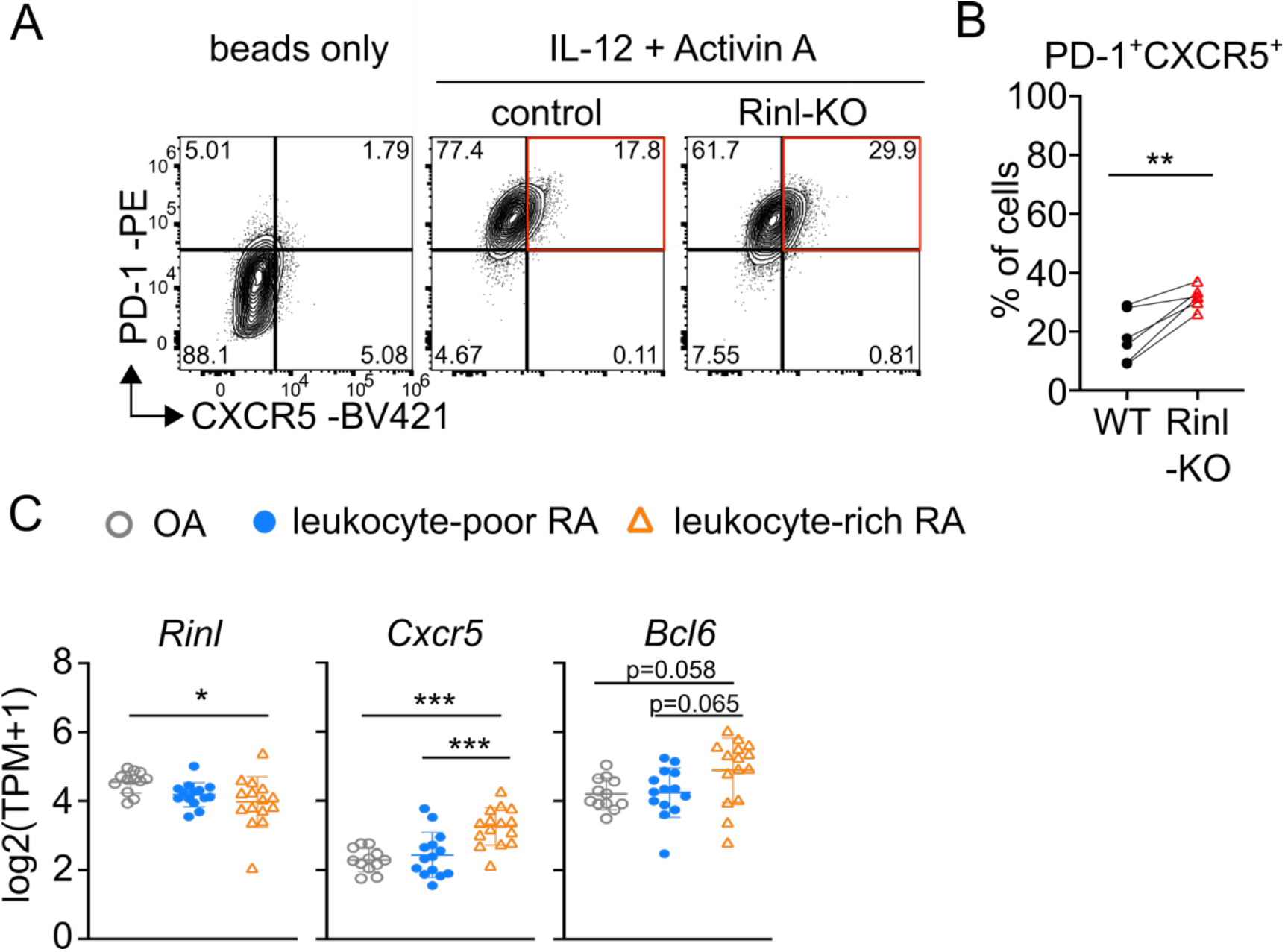
Rinl negatively regulates Tfh differentiation in human CD4^+^ T cells. **(A)** Representative contour plots showing human PD-1^+^CXCR5^+^ Tfh cells. Rinl was knocked-out in human CD4^+^ T cells and control and Rinl-KO cells were subsequently cultured under Tfh-skewing conditions for 5 days. **(B)** Summary of data shown in **(A). (C)** *Rinl, Cxcr5* and *Bcl6* expression in T cells retrieved from synovial biopsies of different patient cohorts (Zhang *et al*, 2019). OA, Osteoarthritis; RA, Rheumatoid arthritis; TPM, transcripts per million. Data show a summary of 6 donors analyzed in 2 independent experiments **(A**,**B)**. Data were statistically analyzed using paired two-tailed t-tests **(B)** or 1-way ANOVA analysis followed by Tukey’s multiple-comparisons test **(C)**. *p < 0.05, ** p<0.01, ***p<0.001.

To further investigate a potential link between Rinl expression, Tfh differentiation and autoimmunity, we determined the expression levels of Rinl in previously published bulk RNAseq data from T cells in synovial biopsies from human osteoarthritis (OA) and rheumatoid arthritis (RA) patients (Zhang et al., 2019). OA and leukocyte-poor RA patients are considered as non or less inflammatory whereas leukocyte-rich RA patients are showing highest inflammation score. Interestingly, we observed reduced levels of *Rinl* in T cells of leukocyte-rich RA patients as compared to non-inflammatory OA cohort while *Cxcr5* expression is highest in leukocyte-rich RA patients **(Fig. 7C)**. These data indicate that Rinl expression might be beneficial in disease outcome of RA as it negatively regulates Tfh differentiation, the driving factor of disease (Ma et al., 2012).

## Discussion

Tfh are crucial components of humoral immunity. To date, numerous positive and negative regulators including transcription factors, surface receptors, cytokines, ubiquitin ligases, and miRNA were identified as critical components during the multifactorial and multistep process of Tfh differentiation (Jogdand et al., 2016). However, scientific efforts still focus on identifying more factors with the aim to harness Tfh for novel therapeutics and vaccine development.

This study is the first report of the Rinl-KO mouse model and we demonstrate that the novel guanine nucleotide exchange factor Rinl is an important determinant of T follicular helper (Tfh) cell differentiation. We identified that Rinl controls Tfh generation under homeostatic conditions, after immunizations with Th2 and Th1/Th17 driving adjuvants, in acute viral infection, during aging, and also in human CD4^+^ T cells. Therefore, Rinl is not regulating Tfh in a particular setting or species but affects Tfh in general. This places Rinl as an universal negative regulator of Tfh cells independently of their subset specification. This is a contrast to other Tfh regulators, as for instance in mice carrying an activated PI3Kδ, Tfh differentiation was enhanced at baseline and in response to immunization, but not in an acute LCMV infection setting (Preite et al., 2019). Moreover, a series of adoptive transfer experiments clearly indicate that Rinl dampens Tfh differentiation in a T-cell intrinsic manner without affecting their *in vivo* expansion or migration ability. Importantly, *in vitro* as well as *in vivo* during immunization studies or viral infection experiments, other T helper subsets, such as Th1, Th2, Th17 or Treg, are not affected in the absence of Rinl, revealing that Rinl acts specifically on Tfh differentiation.

Mechanistically, we identified that Rinl acts via controlling CD28 signaling placing it as a new factor in CD28 biology. Several studies have demonstrated the central role of CD28 co-stimulation in Tfh generation and as a consequence GC establishment and humoral immune response (Ferguson et al., 1996; Lane et al., 1994; Linterman et al., 2014; Weber et al., 2015). We demonstrate that Rinl slows down CD28 internalization and that after short-term and long-term activation of cells with αCD3ε and αCD28, phosphorylation of downstream molecules is altered. After short-term activation, we detect reduced phosphorylation of SLP76 at Ser376 which was shown to negatively regulate T cell receptor signaling (Di Bartolo et al., 2007). The effect is stronger in cells stimulated with αCD3ε +αCD28 compared to cells solely activated with αCD3ε, suggesting that in absence of Rinl TCR signaling was not dampened as strongly by CD28 co-stimulation. It is therefore tempting to hypothesize that Rinl is rewiring CD28 specific signaling pathways which orchestrate Tfh differentiation by acting on CD28 internalization rate.

Following long-term activation of cells with αCD3ε and αCD28 for 48 hours, Erk1/2 phosphorylation was reduced in Rinl-KO CD4^+^ T cells. This is an important observation, as a recent study showed the requirement for suppression of CD28-mediated Erk1/2 activation at a later stage of T cell activation for Tfh differentiation (Wan et al., 2021). CD28 nucleates different signaling pathways which need to be orchestrated at multiple stages and correctly timed to induce a particular T helper specification, and we describe here a previously unrecognized mechanism by which Rinl regulates these events, by acting via an SLP76-dependent pathway early in CD28 signaling and via dampening of Erk1/2 at a later stage, a regulation which matches the multistage differentiation requirements of Tfh cells.

Another important finding of our study is that Rinl has an effect on CD28 signaling strength. Rinl-KO cells displayed increased proliferation and IL-2 production with low anti-CD28 concentrations compared to WT cells but not at high αCD28 concentration. This is of high interest as CD4+ T cells that are IL-2 producers were shown to differentiate into Tfh cells establishing IL-2 production as an early marker for cells fating towards this helper subset (DiToro et al., 2018). Indeed, our observation that there was a tendency of higher *Cxcr5* expression in Rinl-KO cells activated with lower αCD28 concentration suggests that these cells are more likely to differentiate into Tfh. As we were performing our *in vitro* studies without the addition of Tfh-driving cytokines such as IL-6 or IL-21 (Suto et al., 2008), this is a strong indication that in absence of Rinl, CD4^+^ T cells gain a higher Tfh differentiation potential due to alterations in CD28 signaling strength.

In addition, to study whether the alterations in CD28 endocytosis and signaling are relevant in Tfh differentiation *in vivo*, we aimed to rescue our phenotype by providing additional CD28 stimulation. Interestingly, we were indeed able to achieve same frequencies of Tfh after treatment with αCD28 in WT and Rinl-KO mice suggesting that the dysregulation of the CD28 pathway also plays a role *in vivo*. In summary, our mechanistic studies indicate that the regulation of Tfh differentiation by Rinl operates via modulation of CD28 function *in vitro* and *in vivo*.

Importantly, genetic variants at CD28 loci are associated with rheumatoid arthritis (Raychaudhuri et al., 2009), underscoring the importance of the CD28 pathway as gate keeper of proper immune responses. Of note, Rinl had a lower expression in cells from leukocyte rich synovial fluids of RA patients which also showed increased Tfh hallmarks compared to cells from synovial fluids of patients with less inflammatory phenotype. This indicates that Rinl might control the differentiation of Tfh during rheumatoid arthritis and suggests it as a biomarker of this specific disease, opening new directions of investigation. Therefore, our study has also an important implication for therapeutic considerations.

Finally, various studies indicate that angioimmunoblastic T-cell lymphoma (AITL), a subtype of Peripheral T-cell lymphomas (PTCLs) is derived from Tfh (de Leval et al., 2007; Roncador et al., 2007), and AITL is associated with two recurrent mutation sites of CD28 (Rohr et al., 2016). It was proposed that these mutations tune CD28 signaling by affecting CD28 endocytosis. Consequently, CD28 endocytosis might be a central regulatory mechanism how CD4^+^ T cells control Tfh differentiation and further oncogenicity/malignancy. Therefore, our study highlights a novel regulator of Tfh differentiation and in addition a potential novel regulatory mechanism driving T helper fate decision based on CD28 endocytosis. Future studies with CD28 mutants engineered into human CD4^+^ T cells under Tfh differentiation conditions, as well as CD28 endocytosis studies and their consequence on CD28 signaling are warranted.

Taken together, our study provides an in-depth analysis of Rinl function in lymphocyte development, homeostasis and activation. It reveals a previously unrecognized mechanism by which a guanine nucleotide exchange factor controls differentiation of Tfh in various contexts. Our data provide evidence how GEF function might regulate the generation of T helper cells by controlling receptor trafficking in homeostasis and upon activation and may, if dysregulated, lead to disease.

### Limitations of our study

Our study showed how a novel GEF involved in endocytic processes regulates Tfh differentiation *in vitro* and *in vivo* in a T cell-intrinsic way by regulating CD28 endocytosis and signal transduction. Due to the lack of specific antibodies against murine Rinl and technical limitations in visualizing intracellular compartments in murine primary CD4^+^ T cells, our study did not address whether Rinl and CD28 directly interact or co-localize in the same intracellular compartment after endocytosis. The generation of suitable anti-Rinl antibodies for confocal microscopy will be focus of future approaches and will be key for further mechanistic studies.

## Material and Methods

### Mice

Rin-like deficient mice were generated using a targeting construct containing a STOP codon in the reading frame of exon 4. The STOP codon was followed by the IRES-LacZ::GFP cassette and a floxed neomycin. Subsequently, Rinl^+/KI-neo^ mice were crossed with CMV-Cre mice to delete neomycin resulting in Rinl^+/KI^ mice.

OT-II TCR transgenic mice were provided by Michael Bonelli. TCRα KO mice were kindly provided by Iris Gratz.

CD45.1^+^ congenic mice were obtained from the European Mouse Mutant Archive (EM:01998). Rin-like genotyping was performed from toetip DNA via PCR using primers as follows:

5′-TGGAAGATGGGTCCAGCACT-3′; 5′-GCAGCTCCCTTTGCTCTTGA-3′; 5′-GCCACAAGTTCAGCGTGTCC-3′

All animal experimentation protocols were evaluated by the ethics committee of the Medical University of Vienna and approved by the Federal Ministry for Science and Research, Vienna, Austria. Animal husbandry and experimentation was performed according to the Federation of Laboratory Animal Science Associations (FELASA) guidelines under national laws (Federal Ministry for Economy and Science, Vienna, Austria). These guidelines match those of ARRIVE.

Analyzed mice were of mixed sex and between 8-12 weeks old unless otherwise stated.

### LCMV infection

For infection, 2 ×10^5^ PFU LCMV Armstrong strain viruses were injected intraperitoneally and spleens were analyzed 8 or 21 days post infection.

### Immunization

Antigen mixture for immunization was prepared by mixing OVA_(323–339)_ peptide (Sigma-Aldrich) or NP-OVAL (Ovalbumin, Biosearch technologies) in PBS and Imject™ Alum Adjuvant (Thermo Scientific™) while vortexing and afterwards rotation for 2-3 hours at 4°C. Alternatively Complete Freud’s Adjuvant (CFA) was used as adjuvant.

Mice were immunized with 100 μg/mL NP-OVA/Alum interperitoneally (i.p). For flow cytometry analysis spleens were harvested on day 7 after immunization.

Subcutaneous (s.c.) Footpad (FP) immunization was performed with 10μg/mL OVA_(323–339)_/Alum or CFA or 10 μg/mL NP-OVA/Alum in each FP and analysis was done as indicated in figure legends.

For the *in vivo* CD28 study, 100 μg/mL of anti-mouse CD28 (Clone 37,15, BioXcell) or isotype control (isotype poly Syrian hamster IgG, BioXcell) were injected i.p. one day before immunization.

### Adoptive T cell transfer

WT OT-II^+^ CD45.1^+^, WT OT-II^+^ CD45.2^+^ or Rinl-KO OT-II^+^ CD45.2^+^ naïve CD4^+^ T cells were enriched using negative selection with magnetic beads (naïve CD4^+^ T cell isolation kit, Mitlenyi Biotec) according to manufacturer’s protocol. After being washed with PBS, 1*10^5^ – 1*10^6^ naïve (CD62L^+^CD44L^−^) OT-II^+^ CD4^+^ T cells in 100μl PBS were injected retro-orbitally into immobilized recipient mice (CD45.1, CD45.1xCD45.2, TCRα-KO or TCRα-KO Rinl-KO). For analysis of proliferation, cells were labelled with CFSE (Molecular Probes, Eugene, OR) by incubating 1*10^7^ cells/ml in PBS with 10 mM CFSE (Molecular Probes, Eugene, OR) for 10 min at room temperature before injection. Approximately 16 hours after T cell transfer immunization in each FP was performed. Draining lymph nodes (dLN), non-draining lymph nodes (ndLNs) and spleens were analyzed by flow cytometry 2 or 7 days after immunization.

For histology of dLNs, 1*10^6^, 2*10^5^ or 1*10^5^ naïve OT-II^+^ CD4^+^ were transferred and dLNs were collected at days 3, 5 and 7 after immunization, respectively.

### Isolation and activation/differentiation of CD4^+^ T cells

CD4^+^ T cells were isolated from pooled axillary, brachial and inguinal lymph nodes (LN) and spleens of WT and Rinl-KO mice. The cell suspensions were incubated with biotinylated anti-CD8α (RRID: AB_312743; 53-6.7), anti-CD11b (AB_312787; MEL1/70), anti-B220 (AB_312988; RA3-6B2), anti–Gr1 (AB_313369; Ly-6g), anti-Ter119 (AB_313705; Ter-119), anti-CD11c (AB_313773; N418), anti-NK1.1 (AB_313391; PK136) Abs in PBS supplemented with 2% FBS. Antibodies were purchased from BioLegend. Subsequently, CD4^+^ T cells were pre-enriched by magnetic negative depletion using streptavidin beads (MagniSort SAV Negative Selection beads, Thermo Scientific) according to the manufacturer’s description. For total CD4^+^, cells were used immediately. For naïve CD4^+^ T cells, pre-enriched CD4^+^ T cells were stained and sorted for CD25^−^ CD44^low^CD62L^+^ on a SH800 (SONY).

For *in vitro* activation studies, purified naïve CD4^+^ T cells were labelled with CFSE or cell Proliferation Dye eFluor™ 450 (eBioscience™) by incubating 1*10^7^ cells/ml in PBS with 10 mM CFSE for 10 min at room temperature. The labeling reaction was stopped by adding T cell medium. Afterwards, 1*10^5^ cells in 200μl/well complete T-cell medium (RPMI1640 supplemented with 10% FCS (Sigma/Biowest), antibiotics [PenStrep], Glutmax, 50 mM βME) were seeded on 96-well flat-bottom plates. The plates were pre-coated overnight with 1 μg/mL anti-CD3ε (AB_394590; BD Biosciences) and different anti-CD28 (AB_394763; BD Biosciences) concentrations. Cells were harvested and analyzed on day 3. Additionally, supernatant was kept for analysis of IL-2.

Th1/Th17 and iTreg cells were generated from sorted naïve CD4^+^ T cells activated with 1μg/mL anti-CD3ε/ anti-CD28 for 3 days in complete T cell medium supplemented with different cytokine mixes. For Th1 condition, 5ng/mL IL-12 (R&D Systems), 20 U/mL rhIL-2 (PeproTech), and 3μg/mL anti–IL-4 (BioXcell) were used. For Th17 condition 20 ng/mL IL-6, 1 ng/mL rhTGF-β1 (BioLegend), 10 ng/mL IL-1β (BioLegend) and 20 ng/mL IL-23 (R&D) were used and for iTreg condition 1 ng/mL rhTGFβ1. Th2 cells were generated by activating naïve CD4^+^ T cells with 1 μg/mL anti-CD3ε/3 μg/mL anti-CD28 for 3 days in complete T cell medium supplemented with 250U recombinant IL-4 (PeproTech), 6 μg/mL anti-IL-12 (BioXcell), 10 μg/mL anti-IFNγ (BioXcell) and 10 U/mL rhIL-2 (PeproTech). After 3 and 5 days, cells were split and fresh blocking antibodies were added. Cells were analyzed at day 6.

### CRISPR/ Cas9 mediated deletion of Rinl in human CD4^+^ T cells and human Tfh culture

All functional assays were performed in IMDM (Gibco, Thermo Fisher Scientific) supplemented with 10 % of fetal calf serum (Gibco) and 10 μg/mL of gentamycin (Gibco). Peripheral blood draws were performed from healthy human volunteers in accordance with the Ethics Committee of the Medical University of Vienna (EC number EK 1150/2015). Mononuclear cells were isolated by standard Ficoll-Paque centrifugation. Naïve human CD4^+^ T cells were isolated using the EasySep™ Human Naïve CD4^+^ T Cell Isolation Kit II (Stem Cell Technologies) according to the manufacturers’ instructions. Purity of isolated cells was assessed by flow cytometry and found > 95% for all specimen. Subsequently, Crispr/Cas9 knockout of human Rinl was performed as described before (Seki and Rutz, 2018). In detail, 1 μL of a mixture of two Rinl specific crRNAs (Alt-R® CRISPR-Cas9 crRNA; total concentration 320μM; sequences: TGGACCCTGCCGATCTGCAC <u>AGG</u>; TGAATGGTGAGCACGTCCTC <u>AGG</u>; underlined is PAM sequence) were mixed with 1 μL tracr RNA (320μM; all Integrated DNA Technologies) and hybridized for 10min at room temperature. The crRNA-tracrRNA duplex was subsequently complexed with 0,7 μL recombinant Cas9 (Alt-R® S.p. Cas9 Nuclease V; 10 μg/μL; IDT) for 30min at 37°C. Similarly, a control RNP complex was assembled using a non-targeting crRNA (Alt-R® CRISPR-Cas9 Negative Control crRNA #1; IDT). For electroporation, 1*10^6^ purified naïve T-cells were resuspended in 15 μL buffer P3 + 5 μL Supplement 1 (P3 Primary Cell 4D-NucleofectorTM X Kit S; Lonza) and mixed with 2.7 μL RNP in the 16-well strips provided in the kit. Cells were electroporated on a 4D nucleofector (4D-Nucleofector Core Unit, 4D-Nucleofector X Unit; Lonza) using the pulse code EH100. Immediately afterwards, 80 μl pre-warmed fresh medium were added to the cells. After one hour of resting, cells were transferred to 24-well plates and incubated for three days in medium containing 10 U/mL rhIL-2 (Peprotech) to allow establishment of the knockout. Crispr/Cas9 knockout efficiency was determined by Sanger sequencing of the target sites of the two gRNAs and analyzed using the Synthego inference of CRISPR edits analysis tool (ICE v2 CRISPR Analysis Tool; Synthego, Menlo Park, CA) and the knockout score defining frameshift insertions/deletions was found to be > 50 % for both loci in all samples tested.

Human Tfh culture was performed as described previously (Locci et al., 2016).

### Extracellular and intracellular stainings

Thymii, spleens, peripheral lymph nodes and Peyer’s patches were removed and passaged through a 70μm nylon cell strainer to receive single cells suspensions. Erythrocytes were removed by using Pharmlyse (BioLegend). 2*10^6^ cells were incubated with Fc block (BD Pharmingen) and subsequently stained for surface markers with fluorophore-conjugated antibodies for 30 minutes on ice.

For intracellular transcription factor staining, cells were fixed and permeabilized using the FoxP3 staining buffer set (Thermo Scientific) according to manufacturer’s protocol. For intracellular cytokine staining, cells were stimulated with phorbol 12-myristate 13-acetate (PMA, 25 ng/mL) and ionomycin (750 ng/mL; both Sigma-Aldrich) in the presence of GolgiStop (4 μl/6mL; BD Biosciences) for 3-4 hours. Cells were then fixed/permeabilized using Cytofix/Cytoperm (BD Biosciences). Measurements were performed with a BDFortessa and analyzed using FlowJo 10.2 software (TreeStar).

### Phospho staining

Intracellular staining of pSLP76_Ser376_, pSLP76_Y128_, pErk1/2_T202/Y204_ and pAkt_S473_ of activated CD4^+^ T cells was performed by fixing for 15 minutes at 37 °C with BD Cytofix (BD Biosciences), permeabilization for 30min on ice with BD Phosflow™ Perm Buffer III (BD Biosciences) and staining with the indicated phospho antibodies in PBS/2% FCS for 1 hour in the dark at 4°C.

### Short term activation assay

For short term activation, 0,5*10^6^ CD4^+^ T cells were incubated with biotinylated 1μg/mL anti-CD3ε (Clone 145-2C11, BioLegend) alone or in combination with 2ug/mL biotinylated anti-CD28 (Clone 37,15, BioLegend) for 15min at room temperature. Streptavidin (1ug/mL) was added and samples were transferred to 37°C for 2 minutes. The reaction was stopped by addition of Cytofix and subsequent staining with phosphor-antibodies.

### Confocal microscopy of dLNs

Draining LNs were harvested and fixed in Antigenfix (Diapath) for 1,5 hours, washed with PBS and then dehydrated in 30% sucrose prior to embedding in OCT freezing media (Bio-Optica). 20 μm sections were cut on a CM1520 cryostat (Leica) and adhered to Superfrost Plus slides (Thermo Fisher Scientific). Sections were then blocked in PBS containing 0.3% Triton X-100 (Sigma-Aldrich) and 0.5% BSA followed by incubation with Fc block in the same blocking buffer. The following primary Abs were used for staining: BV421-conjugated anti-CD45R/B220 (RA3-6B2, BioLegend), AF488-conjugated anti-CD45.2 (104, BioLegend), biotinylated anti-LYVE-1 (ALY7, eBiosciences), BV421-conjugated anti-TCRβ (H57-597, BioLegend), biotinylated CD21/35 (7E9, Biolegend). The following secondary Abs were used for staining: Streptavidin, Alexa Fluor™ 555 or Alexa Fluor™ 647 Conjugate (Invitrogen). Images were acquired on an inverted Leica microscope (TCS STED CW SP8, Leica Microsystems). A motorized stage was used for tiled imaging. Image analysis was performed using Imaris Microscopy Image Analysis Software (Version9.1.2, Oxford Instruments).

### Internalization assay

Internalization assay was performed as previously described with minor changes (Finetti et al., 2014). Shortly, 2*10^6^ CD4^+^ T cells were seeded out in 96-well round-bottom plates (Sarstedt) and incubated with 1 μg/mL biotinylated anti-CD28 antibody (Clone 37.15, BioLegend) PBS+2%FCS for 30 minutes on ice to allow binding. After washing steps, cells were incubated at 37°C to induce endocytosis. 30uL cell suspension were transferred immediately into a new plate on ice (time point=0). Subsequently, aliquots were collected at different time points and internalization was stopped by incubating the cells on ice. Samples were analyzed by detecting residual surface molecules by staining with an fluorochrome-labelled streptavidin antibody. Samples were handled in duplicates or triplicates. Measurements were performed with a BDFortessa.

### BMDC generation/ co-culture

BMDCs were generated according to published protocols (Madaan et al., 2014). Briefly, bone marrow was isolated from tibiae and femurs and single-cell suspensions were cultured at 10*10^6^ cells per 100 mm bacteriological Falcon petri dish in 10 ml complete T cell medium supplemented with 20 ng/ml GM-CSF (PeproTech, Rocky Hill, NJ). At day 3, additional 10mL complete T cell medium+ 20ng/mL GM-CSF were added. At days 6 and 9, half of the medium was removed, cells were centrifuged and added with freshly prepared 10mL complete T-cell medium+ 20 ng/mL GM-CSF. Non-adherent immature BMDCs were harvested on day 10 and stimulated with OVA_(323–339)_ peptide (1 μg/mL) for 45 minutes at 37°C. Afterwards, they were cultured at a 1:10 ratio with sorted naïve CD4^+^ OT-II^+^ T cells. At day 3, T cells were harvested and analyzed. For assessment of proliferation, T cells were labeled using Cell Proliferation Dye eFluor®450 (Thermo Fisher Scientific) according to manufacturer’s protocol prior to the co-culture. To study the effect of CD28 blocking on proliferation, 40 μg/mL CTLA4-Ig were added to the culture.

### Semiquantitative RT-PCR

Total RNA was isolated total organs or from sorted B220^+^ B cells, GC B cells, CD4^+^/CD8^+^ T cells and Tfh from indicated organs using RNeasy Mini Kit (Qiagen Inc.). cDNA synthesis was performed using Superscript III First Strand Synthesis System (Invitrogen).

### RT-qPCR

Total RNA was extracted from *in vitro* activated CD4^+^ T cells using RNeasy Mini Kit (Qiagen Inc.) as described in manufacturer’s protocol including an on-column digestion step (RNaseFree DNase Set, Qiagen Inc.). RNA was reverse transcribed with Superscript III First Strand Synthesis System (Invitrogen) as described in manufacturer’s protocol. Quantitative real-time PCR (qRTPCR) analysis was performed with the iTaq™ Universal SYBR Green Supermix (Bio-Rad) on the CFX 96 Real-Time PCR detection system (Bio-Rad). The following primers were used: *Cxcr5* fw: ATGAACTACCCACTAACCCTGG. *Cxcr5* rev: TGTAGGGGAATCTCCGTGCT (Wan et al., 2021); *Hprt* fw: GATACAGGCCAGACTTTGTTG; *Hprt* rev: GGTAGGCTGGCCTATAGGCT (Bilic et al., 2006). All experiments were performed in duplicates or triplicates and normalized to the reference gene *Hprt*.

### IL-2 ELISA

IL-2 ELISA was performed using an ELISA MAX™ Deluxe Set Mouse IL-2 as described in manufacturer’s protocol (BioLegend). The O.D. was measured with an ELISA Reader (Multiscan GO, SkanIt Software for Microplate Readers RE) at 450nM.

### LCMV-specific IgG1 ELISA

For measurement of LCMV-specific IgG1 antibodies, ELISA were performed as previously described with minor adaptations (Baumjohann et al., 2013). Briefly, 96-well Nunc MaxiSorp™ flat-bottom plates (Thermo Scientific) were coated with lysates of LCMV Armstrong-infected baby hamster kidney (BHK) cells overnight. Nonspecific binding was blocked with PBS+ 0.5% Tween® 20 (Promega) + 10% FBS for 2 hours at RT. Subsequently, serially diluted serum was added and incubated another 2 hours at RT. Rat anti-mouse IgG1 (Clone A85-1, BD Biosciences) and ECL anti-rat IgG Horseradish Peroxidase linked whole Ab from goat (GE Healthcare UK Limited) were used to detect IgG1 antibodies upon addition of TMB substrate (TMB substrate set, Biolegend). The O.D. was measured with an ELISA Reader (Multiscan GO, SkanIt Software for Microplate Readers RE) at 450nM.

### Bulk RNA-Seq, sample preparation and bioinformatic analysis

Total RNA was prepared from 1-2*10^6^ sorted naïve (CD25^−^ CD44^low^CD62^high^) or effector (CD25^−^ CD44^high^CD62^low^) CD4^+^ T-cells using RNeasy Mini Kit (Qiagen Inc.) as described in manufacturer’s protocol including an on-column digestion step (RNaseFree DNase Set, Qiagen Inc.). The amount of total RNA was quantified using the Qubit 2.0 Fluorometric Quantitation system (Thermo Fisher Scientific) and the RNA integrity number (RIN) was estimated using the Experion Automated Electrophoresis System (Bio-Rad). RNA-Seq libraries were generated by the Biomedical Sequencing facility at CeMM using a Sciclone NGS Workstation (PerkinElmer) and a Zepyhr NGS Workstation (PerkinElmer) with the TruSeq Stranded mRNA LT sample preparation kit (Illumina). Library concentrations were determined with the Qubit 2.0 Fluorometric Quantitation system (Life Technologies, Carlsbad, CA, USA) and were sequenced using the HiSeq 3000/4000 platform (Illumina) following the 50-bp single-read configuration. Following the raw data acquisition (HiSeq Control Software) and base calling (Real-Time Analysis Software, RTA, v2.7.7) that was performed on-instrument, the subsequent raw data processing involved programs based on Picard tools (https://broadinstitute.github.io/picard/) to generate sample-specific, unaligned BAM files. Sequence reads were subsequently mapped onto the mouse reference genome assembly build mm10 (a flavor of GRCm38) using Spliced Transcripts Alignment to a Reference (STAR) aligner (Dobin et al., 2013) which was run with options recommended by the ENCODE project. Aligned reads that overlapped Ensembl transcript features were counted with the Bioconductor (v3.12) GenomicAlignments (v1.26.0) package. Differential expression was tested using the Bioconductor DESeq2 package (v1.30.0)(Love et al., 2014). The volcano plot was generated using GraphPad Prism and downstream analysis was performed using Ingenuity Pathway Analysis (Qiagen IPA).

### Bioinformatics analysis of published RNA-sequencing data

RNA-seq from previously published data (Zhang et al., 2019) was downloaded from Immport (SDY998) as normalized log2 transformed data. Retrieved data was checked for quality and consistency after combining with corresponding metadata. All data manipulation was performed in R (v4.1.2).

### Statistical analysis

All statistical analyses were performed using GraphPadPrism software (version 8.0.2). For single comparisons paired or unpaired Student’s t-test or one-sample T-test were performed as indicated in figure legends. For multiple comparisons, 1-way or 2-way ANOVA followed by Tukey’s multiple-comparisons tests were performed. Horizontal lines indicate the mean and error bars show the standard error of the mean (SEM).

## Supporting information

All supplementary figures for the manuscript

## Data availability statement

The authors declare that the data supporting the findings of this research are available in the article, the Supplementary Material, or on request from the corresponding author.

## Ethics statement

The animal study was reviewed and approved by the Austrian Federal Ministry for Education, Science and Art. Peripheral blood draws were performed from healthy human volunteers in accordance with the Ethics Committee of the Medical University of Vienna (EC number EK 1150/2015)

## Author contributions

LS, RH and NB designed the research; LS performed most of the experiments and analyzed the data. MA, CZ, RR, and PH contributed to *in vivo* experiments; AS performed semiquantitative PCR analysis; SS contributed to LCMV infection experiments and analyzed data. LS, MM, AT and MB performed bioinformatics analysis; BW, RH and IT generated the Rinl-KO mice; ES, MK and MI performed confocal microscopy experiments; TF and KS performed CRISPR/ Cas9 mediated deletion in human CD4^+^ T cells; LS and NB wrote the manuscript with contributions from all co-authors.

## Acknowledgements

We thank the Biomedical Sequencing Facility at CeMM for assistance with next-generation sequencing and Michael Schuster for initial data processing and analysis of RNA-seq data. Confocal immunofluorescence microscopy was carried out at Alembic, San Raffaele Scientific Institute and the Vita-Salute San Raffaele University. Mice were kept at the Core facility laboratory animal breeding and husbandry of the Medical University of Vienna. We also thank Dieter Printz of the FACS Core Unit at St. Anna CCRI for cell sorting and the NIH Tetramer Facility for providing GP66-specific MHC class II tetramers. LS would like to thank the European Federation of Immunology (EFIS) for awarding the EFIS-IL Short-term Fellowship.

## Funding

This study has been funded by Austrian Science Foundation (FWF) through projects P24265, P30885 and F7004. Studies in the lab of S.S. were funded by Austrian Science Fund (FWF) P27747. LS received an EFIS-IL Short-term Fellowship for a research stay at Vita San Raffaele University and Division of Immunology, Transplantation and Infectious Diseases, IRCCS San Raffaele Scientific Institute, Milan, Italy.

## Declaration of interest

The authors declare no competing interests.

## Notes

### Competing Interest Statement

The authors have declared no competing interest.

