## Supplementary material for "The guanine nucleotide exchange factor Rin-like acts as a gatekeeper for T follicular helper cell differentiation via regulating CD28 signaling": All supplementary figures for the manuscript

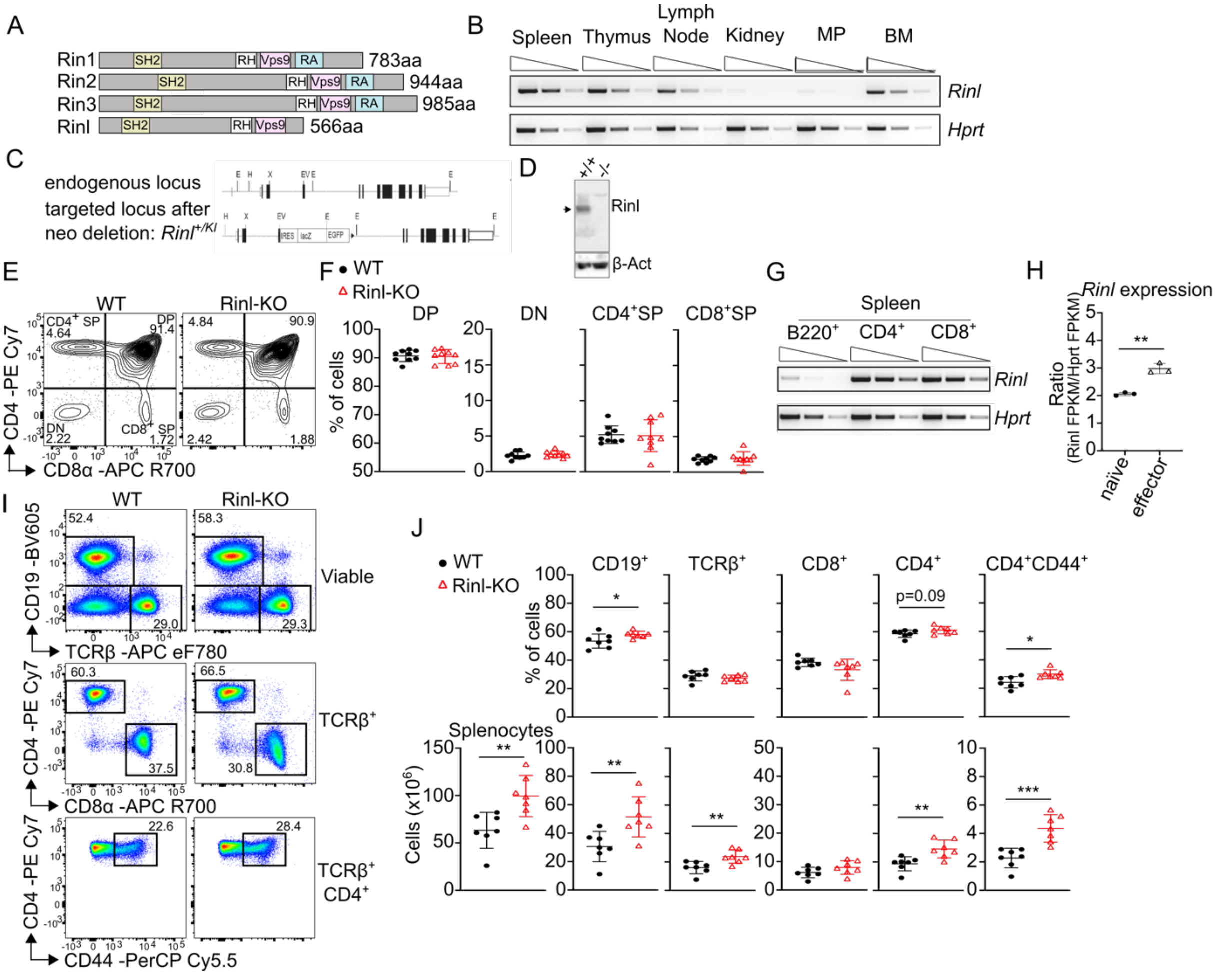


**Supplementary Fig.1: Rinl is highly expressed in lymphoid organs and impacts T cell homeostasis in spleen. (A)** Schematic representation of Ras interaction/interference (Rin) family members Rin1-Rin3 and Rinl. **(B)** Semi-quantitative PCR of *Rinl* expression in different organs of WT mice. *Hprt* expression was used as control. **(C)** Rinl locus before and after targeting. Rinl has 12 exons (depicted as filled black boxes) and exons 1-11 encode for Rinl protein. Introns are represented by connecting lines. A STOP codon in the reading frame of exon 4 followed by IRES-LacZ::GFP cassette and a floxed neomycin was used to inactivate Rinl. Rinl+/KI mice were generated by crossing Rinl+/KI-neo mice with CMV-Cre mice to delete neomycin. **(D)** Immunoblot analysis depicting Rinl expression in thymocytes of the indicated genotype. The arrow indicates the position of Rinl at 60 kDa. β-Actin protein abundance was used to confirm equal loading. **(E)** Representative contour plots of thymocyte gating. **(F)** Summary graphs show the percentages of double positive (DP), double negative (DN), single positive (SP) $CD4^+$ and SP $CD8^+$ thymocytes. **(G)** Semi-quantitative PCR of *Rinl* expression in $B220^+$, $CD4^+$ and $CD8^+$ cells of WT spleens. **(H)** Rinl expression level among naïve ($CD62L^+CD44^-$) and effector ($CD62L^-CD44^+$) $CD4^+$ T cells. Data are expressed as Fragments Per Kilobase of transcript per Million fragments mapped (FPKM) and Rinl expression was normalized to Hprt expression. **(I)** Representative pseudocolour plots for gating of $CD19^+$, $TCR\beta^+$, $CD4^+$, $CD8^+$ and $CD4^+CD44^+$ cells from spleen. **(J)** Quantification (upper panel) and summary of cell numbers (lower panel) of **(I)**. The summary of 9 **(F)** or 7 **(J)** mice per genotype analyzed in 3 independent experiments is shown. Data were statistically analyzed using unpaired two-tailed t-tests. *p < 0.05, **p<0.01, ***p<0.001. Aa, amino acid; SH, Src-homology; RH, Rin homology; Vps, Vacuolar protein sorting-associated protein; RA, Ras association, BM, bone marrow; E, EcoRI; X, XhoI; V; EcoR; H, HindII.

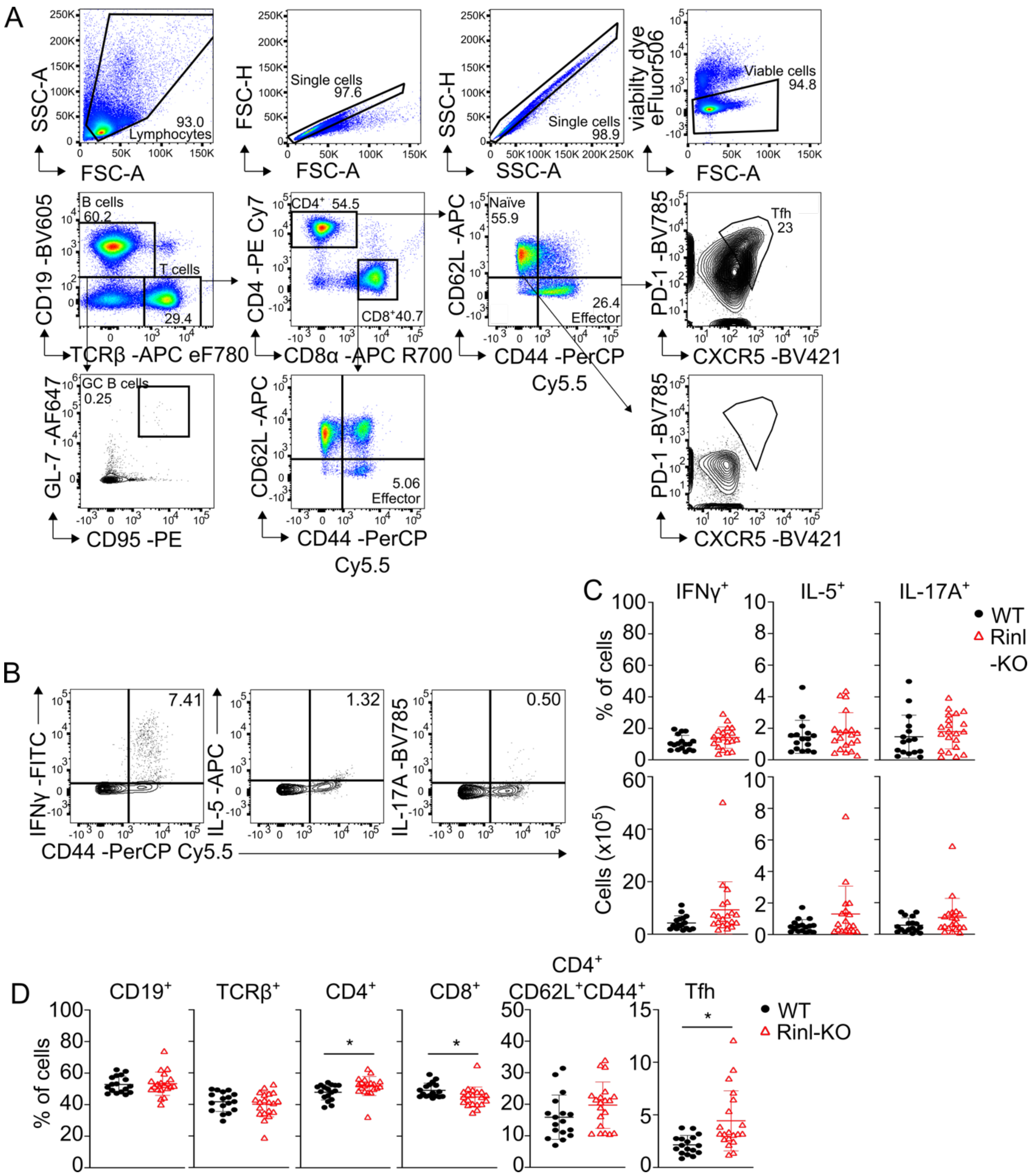


**Supplementary Fig. 2: Loss of Rinl affects peripheral T cell homeostasis in older mice.** **(A)** Gating strategy for analysis of lymphocyte subsets in spleen of older WT and Rinl-KO mice. **(B)** Representative gating of cytokine producing CD4$^{+}$ T cells of spleens of older mice **(C)** Summary graphs of cytokine producing cells. **(D)** Summary graphs of cell subsets in peripheral lymph nodes of older WT and Rinl-KO mice. Summary of 16-20 mice that were analyzed in 3 independent experiments are shown **(C,D).** Data were statistically analyzed using unpaired two-tailed t-tests. *$p < 0.05$.

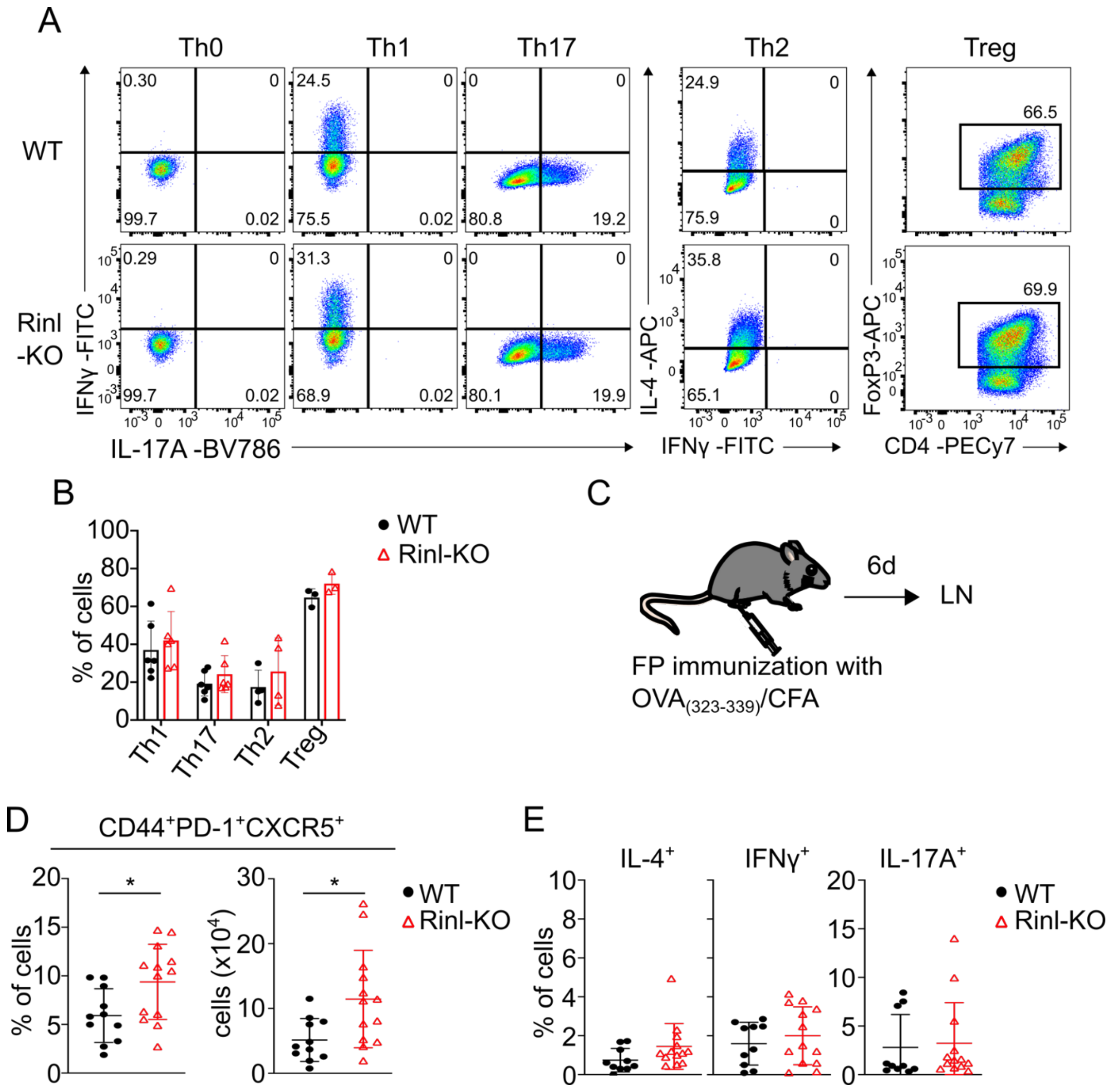


**Supplementary Fig. 3.: Rinl does not regulate T helper cell differentiation *in vitro* and *in vivo*** **(A)** Representative contour plots of cytokine-producing CD4$^{+}$ T cells after *in vitro* differentiation under Th1, Th2, Th17 or Treg skewing conditions. **(B)** Summary of **(A)**. **(C)** Experimental scheme. **(D)** Summary graphs of frequencies and cell numbers of Tfh, **(E)** Summary graphs of cytokine-producing cells. Data show summary of 3-6 **(B)** and 9-13 **(D,E)** mice analyzed in 3-4 independent experiments. Data were statistically analyzed using paired two-tailed t-tests. *$p < 0.05$.

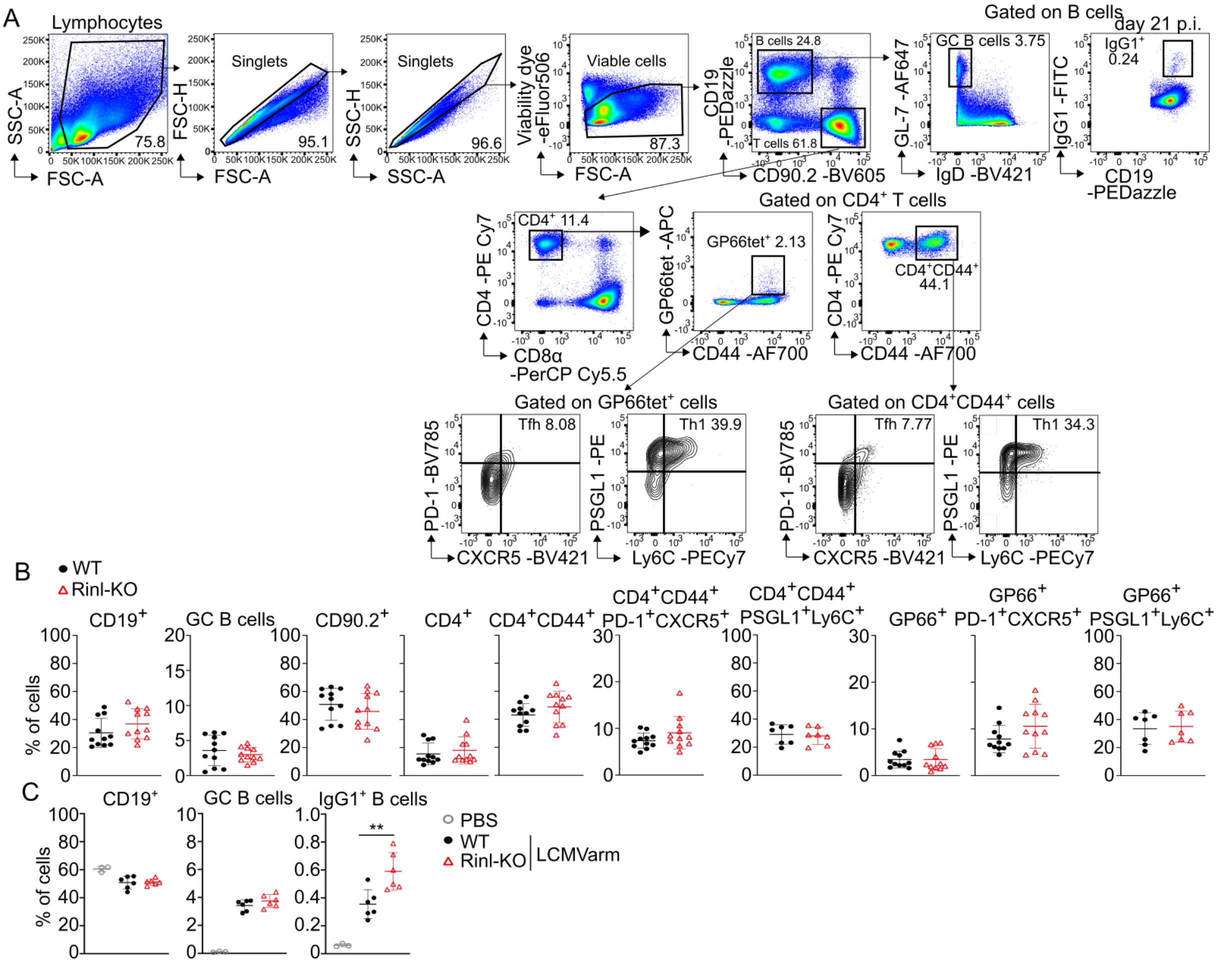


**Supplementary Fig. 4: Analysis of spleens from WT and Rinl-KO mice after LCMV Armstrong infection. (A)** Gating strategy of spleen of WT and Rinl-KO mice after LCMV infection **(B)**. Summary diagrams depict frequencies of lymphocyte subsets from spleens from WT and Rinl-KO mice 8 days post infection. **(C)** Summary diagrams depict frequencies of B cell subsets from spleens from WT and Rinl-KO mice 21 days post infection. Data show a summary of 11 **(B)** or 6 **(C)** mice per group analyzed in 3 **(B)** or 2 **(C)** independent experiments. Data were statistically analyzed using unpaired two-tailed t-tests. **p < 0.01.

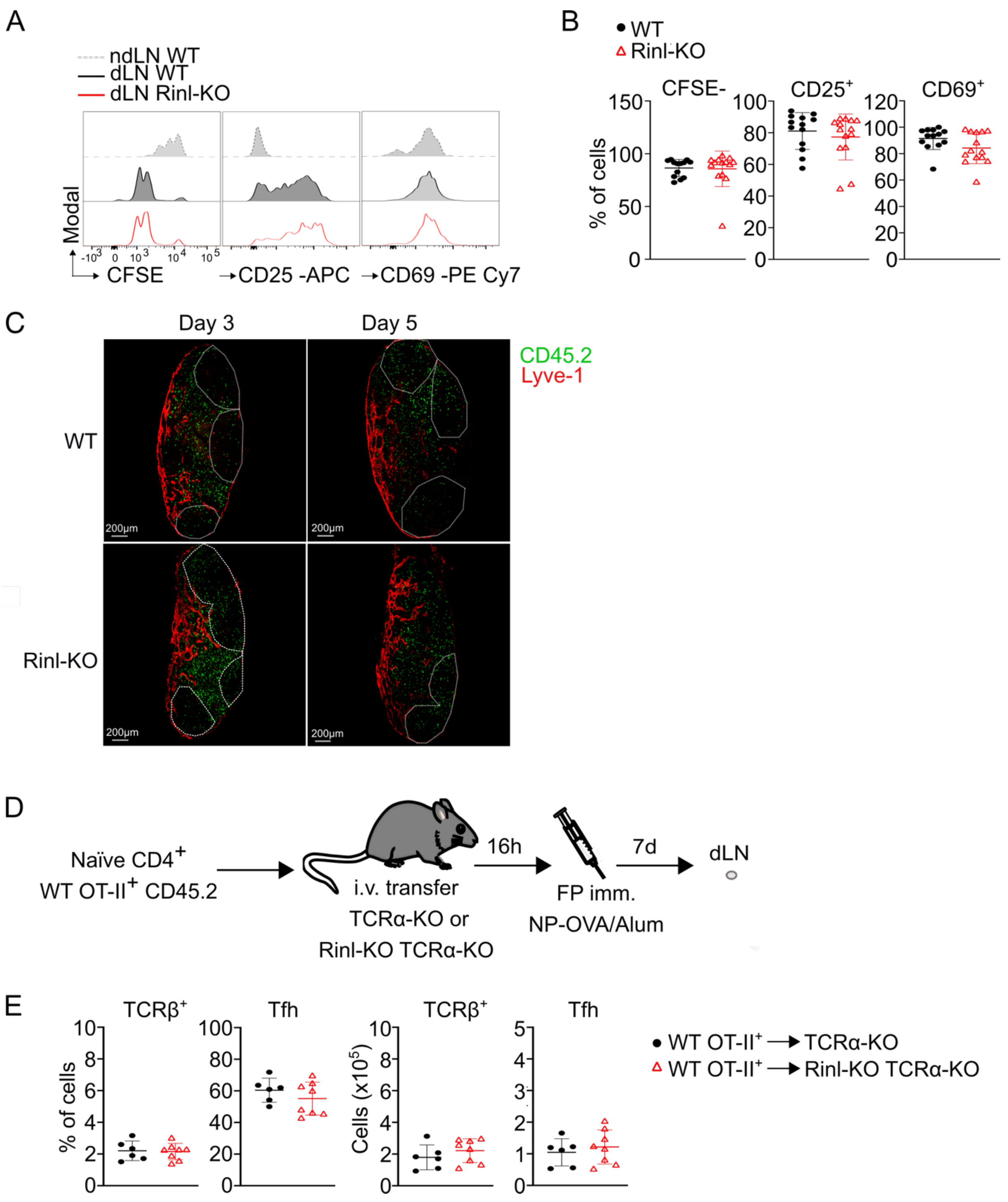


**Supplementary Fig. 5.: Rinl does not control early T cell proliferation and migration *in vivo*. (A)** Representative histograms of proliferation (CFSE) and CD25 and CD69 expression on transferred ($CD45.2^{+}CD45.1^{-}$) $CD4^{+}$ T cells. **(B)** Summary of **(A). (C)** Representative confocal images of dLNs at day 3 and 5 after immunization. Dashed lines represent edges of B cell follicles and were set based on B220 staining. Ag-specific $CD45.2^{+}$ cells are shown in green, Lyve-1 staining in red. Images are representative of at least 3-5 LNs from 2 independent experiments. **(D)** Experimental scheme. **(E)** Summary graphs show percentages and cell counts of TCRβ$^+$ cells and Tfh in TCRα$^{-/-}$ or Rinl$^{-/-}$ TCRα$^{-/-}$ mice. Summary of 13-14 **(B)** and 6-8 **(E)** mice that were analyzed in 3 **(B)** and 2 **(E)** independent experiments are shown**.**

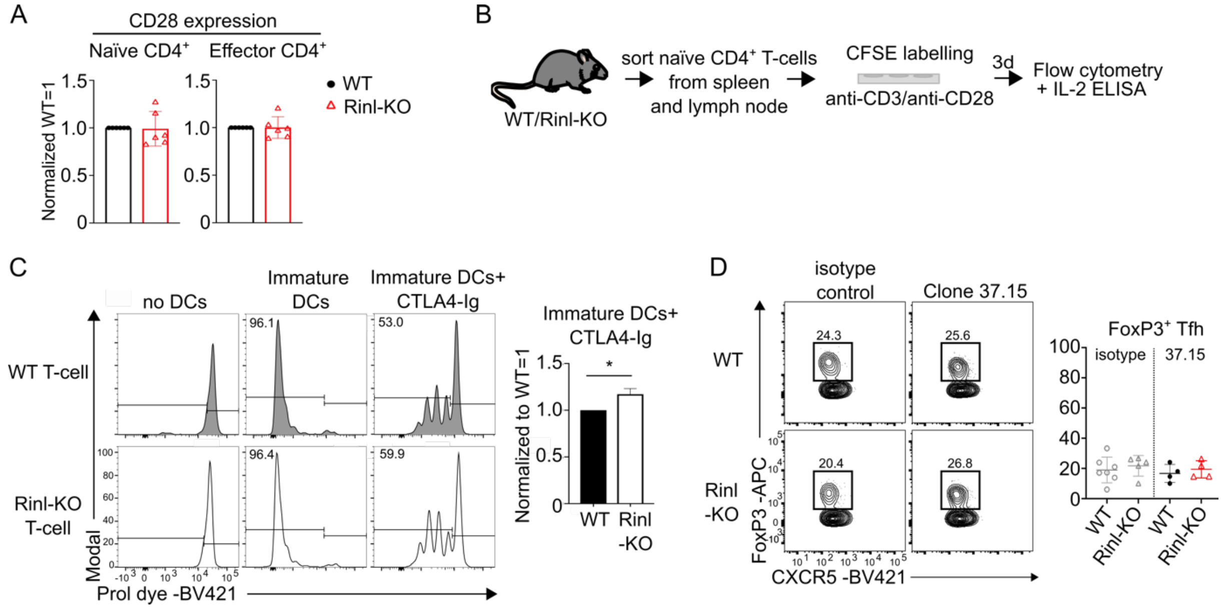


**Supplementary Fig. 6: Rinl regulates CD28 response in CD4$^+$ T cells. (A)** Summary graph shows CD28 expression. **(B)** Experimental scheme for *in vitro* activation. **(C)** Representative histograms depicting proliferation of T cells after activation with bone-marrow derived dendritic cells (BMDCs) ± CTLA4-Ig. Summary is shown alongside **(D)** Representative contour plots of follicular T regulatory cells (Tfr) of WT and Rinl-KO mice treated with isotype control or αCD28 (Clone 37.15) one day before immunization. Analysis is shown alongside. Data show a summary 6 **(A)**, 3 **(C)** or 4-7 **(D)** mice per group that were analyzed in 2 **(D)** or 3 **(A,C)** independent experiments. Data were statistically analyzed using paired two-tailed t-tests. *$p < 0.05$.
